# Single cell profiling of bone metastasis ecosystems from multiple cancer types reveals convergent and divergent mechanisms of bone colonization

**DOI:** 10.1101/2024.05.07.593027

**Authors:** Fengshuo Liu, Yunfeng Ding, Zhan Xu, Xiaoxin Hao, Tianhong Pan, George Miles, Yi-Hsuan Wu, Jun Liu, Igor L. Bado, Weijie Zhang, Ling Wu, Yang Gao, Liqun Yu, David G. Edwards, Hilda L. Chan, Sergio Aguirre, Michael Warren Dieffenbach, Elina Chen, Yichao Shen, Dane Hoffman, Luis Becerra Dominguez, Charlotte Helena Rivas, Xiang Chen, Hai Wang, Zbigniew Gugala, Robert L. Satcher, Xiang H.-F. Zhang

**Affiliations:** Lester and Sue Smith Breast Center, Baylor College of Medicine, One Baylor Plaza, Houston, TX 77030, USA; Dan L. Duncan Cancer Center, Baylor College of Medicine, One Baylor Plaza, Houston, TX 77030, USA; Department of Molecular and Cellular Biology, Baylor College of Medicine, One Baylor Plaza, Houston, TX 77030, USA; Medical Scientist Training Program, Baylor College of Medicine, Houston, TX 77030, USA; Graduate Program in Cancer and Cell Biology, Baylor College of Medicine, One Baylor Plaza, Houston, TX 77030, USA; Graduate Program in Integrative Molecular and Biomedical Sciences, Baylor College of Medicine, One Baylor Plaza, Houston, TX 77030, USA; Graduate Program in Immunology and Microbiology, Baylor College of Medicine, One Baylor Plaza, Houston, TX 77030, USA; Graduate Program in Development, Disease Models, and Therapeutics, Baylor College of Medicine, One Baylor Plaza, Houston TX 77030, USA; Department of Orthopedic Oncology, University of Texas, MD Anderson Cancer Center, Houston, TX, United States of America; Department of Orthopedic Surgery and Rehabilitation, University of Texas Medical Branch, Galveston, TX, USA; McNair Medical Institute, Baylor College of Medicine, One Baylor Plaza, Houston, TX 77030, USA; College of Natural Sciences, University of Texas at Austin, 110 Inner Campus Drive, Austin, TX 78706, USA

## Abstract

Bone represents congenial soil for metastatic seeds and is frequently affected by metastasis of multiple cancer types. The histological and molecular characteristics of bone metastases (BMs) are diverse but poorly understood. Herein, we performed single-cell RNA-seq on 34 BMs from 6 cancer types and identified 3 ecosystem archetypes characterized by enrichment of macrophages/osteoclasts (Mφ-OC), regulatory/exhausted T cells (Treg-Tex), and monocytes (Mono), respectively. Breast cancer BMs are mostly the Mφ-OC archetype driven by the osteolytic vicious cycle, whereas kidney cancers BMs mainly belong to the Treg-Tex archetype that lacks osteoclasts. Lung cancers BMs evenly distributed across all archetypes. Further analyses revealed parallel mechanisms of immunosuppression and bone remodeling. Elevated estrogen signaling distinguishes macrophages in the Mφ-OC subtype, which was investigated in a companion study. Together, we elucidated that divergent mechanisms toward bone colonization and that BMs of different origins can adopt the same mechanism through convergent evolution or adaptation.

**HIGHLIGHTS:**

- Analyses of bone metastases from 6 cancer types revealed three immune archetypes.
- Archetypes diverge on immune trajectories, and features of tumor and stromal cells.
- Dominant cell type in each archetype undergoes convergent evolution.
- Regulatory networks converge on osteoclasts and Tregs to drive archetype formation.

**Graphic Abstract:** 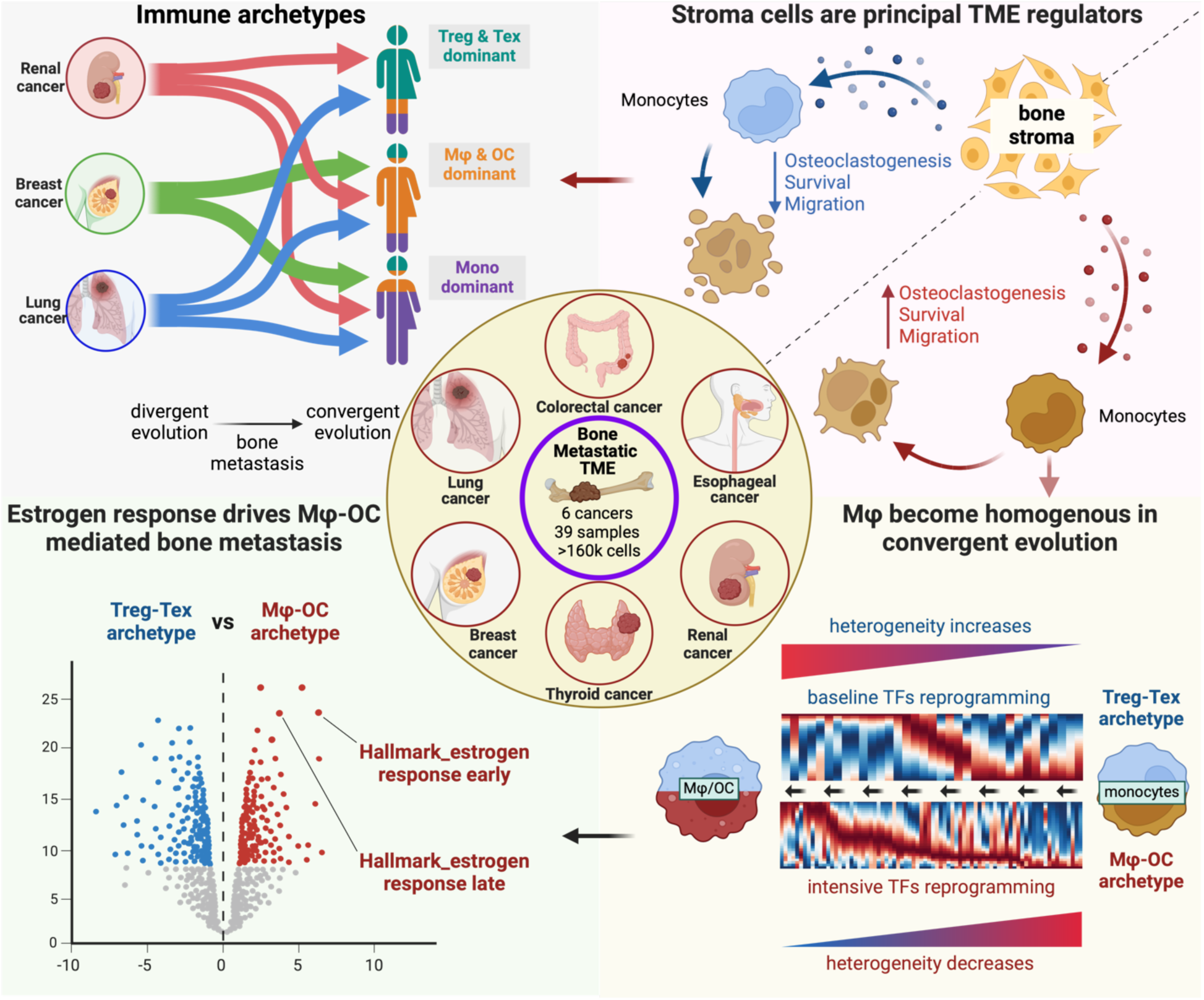

## INTRODUCTION

Bone and bone marrow play multiple roles in normal physiology^1^. The mineralized part of bone provides mechanistic support to the body, whereas the semifluid bone marrow is the birthplace of new blood and immune cells. In addition, bone modeling generates metabolic and endocrine impact that extends to the entire organism^2,3^. The enrichment of cytokines and growth factors in the bone microenvironment provides congenial soil for not only normal stem cells but also metastatic seeds^4^.

Indeed, bone is a common metastatic target of many solid cancer types. Upon arrival, the disseminated tumor cells (DTCs) often reside in the perivascular niche, and their interaction with endothelial cells sometimes enforce a dormant state and sometimes drives mesenchymal-to-epithelial transition and proliferation^5–7^. On the other hand, the outgrowth of DTCs was observed associated with an osteogenic environment wherein cancer cells and osteogenic cells form intimate interactions mediated by heterotypic adherens and gap junctions^8,9^. The transition of DTCs from the perivascular niche to the osteogenic niche may be coupled with the bone remodeling process^10^. This previous knowledge highlights the importance of the crosstalk among metastatic cells, endothelial cells and mesenchymal/osteogenic cells in early-stage bone metastasis.

To fully colonize bone, tumor cells evolve mechanisms to invade the mineralized bone matrix and evade immunosurveillance. Previous studies indicate that dysregulation of osteoclast and osteoblast activities, especially the hyperactive bone resorption driven by osteoclasts, underpin the metastasis-induced remodeling to bone^11–14^. Accordingly, agents targeting osteoclasts, such as bisphosphonates and denosumab (anti-RANKL) have been widely used to treat bone metastases of various cancer types, including osteosclerotic prostate cancers^15^. However, some cancers do not respond to these agents. Renal cancers, for instance, are notoriously resistant to bisphosphonates despite the similar osteolytic appearance^16^, suggesting potentially osteoclast-independent mechanisms.

In addition to the ability to invade the foreign environment, exacerbated immunosuppression is another hallmark of metastases. Compared to visceral and brain metastases, bone metastasis is much less characterized due to the scarcity of clinical specimens^17,18^. The immune landscape of bone metastasis is expected to distinctive considering its unique functions in generating immune cells and its enrichment of adult stem cells, which are postulated to reside in immunopreviliged niches^19^. However, very little is known about the potential immunosuppressive mechanisms in bone metastases.

Prompted by these significant gaps in our knowledge, we have performed single cell RNA-seq (scRNA-seq) of 34 human bone metastases from 6 cancer types. Interestingly, we identified three archetypes of ecosystems that do not completely coincide with organs of origin. We further characterized two of these archetypes represented by the majority of breast cancer and kidney cancer, respectively, and revealed distinctive parallel mechanisms of bone colonization and immune-evasion. Our work demonstrates how different cancer types may colonize the same organ via a finite number of distinct mechanisms, and provides rationale to investigate completely different treatment strategy for certain bone metastases.

## RESULTS

### Human bone metastases of different cancer types fall into three ecosystem archetypes with largely distinctive immune landscape

Fresh bone metastasis tissues were collected during orthopedic surgeries. The clinical information of all 34 patients, including prior treatments, were provided in Supplementary Information. For each sample, a portion was immediately dissociated and FACS sorted into single cell suspensions for scRNA-seq, and the remainder was preserved for other purposes. We used the cell hashing approach to combine 3-5 samples in each run of sequencing to reduce the cost and batch effects. Detailed information of data pre-procession is provided in the Methods section and summarized in **Figure 1A**. H&E-stained tissue sections and accompanying pathology descriptions for representative samples are available for download in the supplementary files.

**Figure 1:**
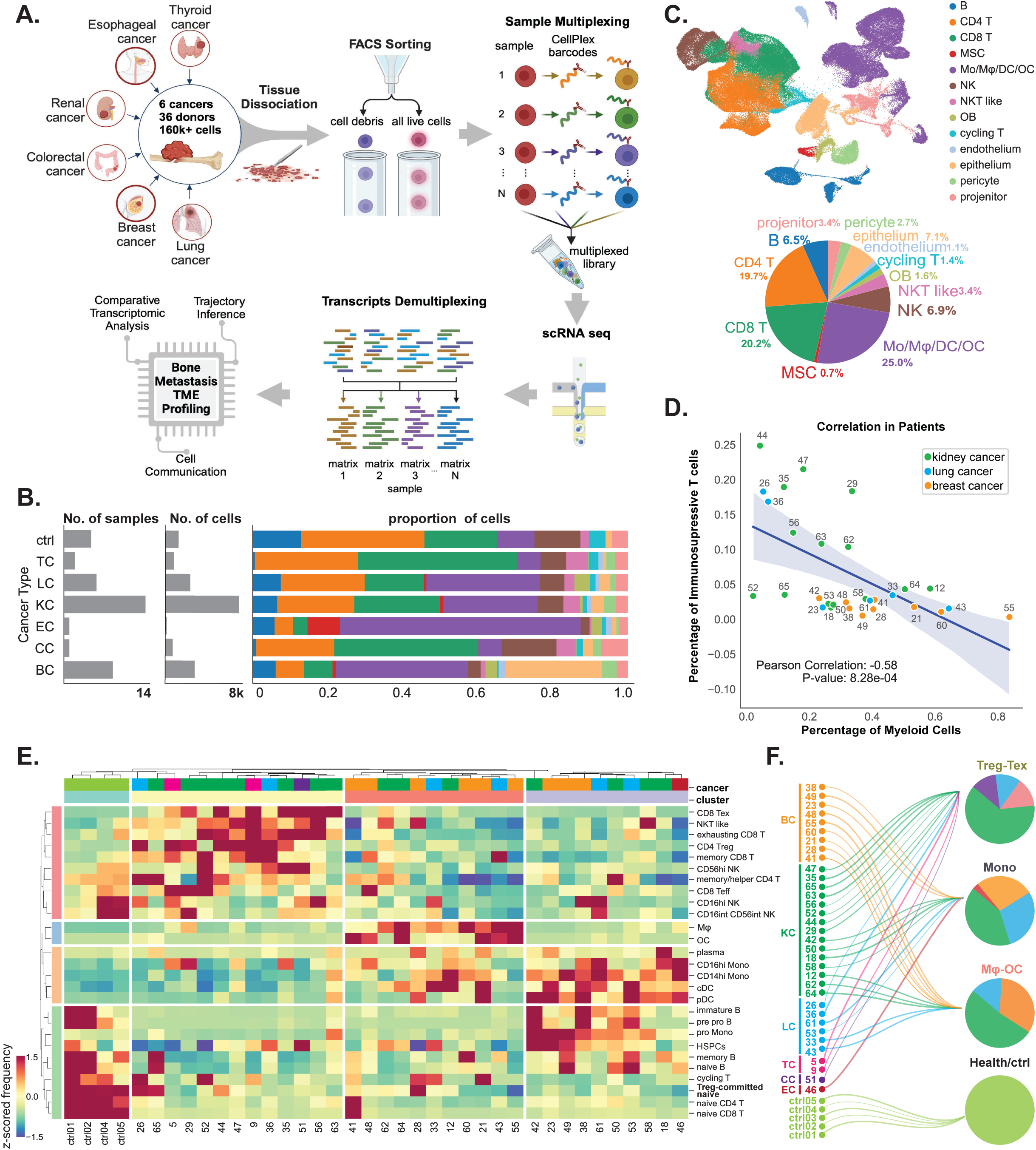
Overview of Study Design and Immune Archetypes Among Patients. **A.** Overview of the study workflow, illustrating the process from data collection to analysis. A total of 39 patients and healthy donors were included, resulting in over 160,000 high-quality cells post-preprocessing and quality control (QC). (Illustration created with BioRender) **B.** Samples and cell count, categorized by cancer types. **C.** UMAP projection depicting major cell types, annotated for clarity. The accompanying pie chart details the proportion of each cell type relative to the overall cell count. Bar graphs represent cell type distributions within individual cancer categories as shown in panel B. **D.** Patient hierarchical clustering based on normalized (z-score standardized across patient data) cell frequencies, revealing four distinct immune archetypes: healthy control donors, monocyte-enriched (Mono), macrophage/osteoclast-enriched (Mφ-OC), and regulatory/exhausted T cell-enriched (Treg-Tex). **E.** Pearson correlation analysis of myeloid cell frequency against the frequency of immunosuppressive T cells (Tregs and Tex) in breast cancer (BC), thyroid cancer (TC), and lung cancer (LC) patients, revealing a significant inverse relationship.

To identify cell types, we assembled a reference cell database based on multiple previously published datasets (see Methods for details) (**Figure S1A**), and used support vector machine (SVM) to classify cells derived from bone metastases. This approach resulted in robust assignment of each cluster of cells to specific cell types (**Figure S1B-1C**), resulting in annotation of 32 cell types (**Figure S1D**). We then manually inspected renowned marker genes of major cell types visually by feature plots (**Figure S1E**) and numerically by gene expression scores (**Figure S1F**), which largely validated SVM annotation.

In bone metastases of all cancer types we observed that immune cells comprise the majority of cellular constituents (**Figure 1B-1C**). This may not be surprising considering that the bone marrow is the birthplace of immune cells. In contrast to bone metastases, previous studies suggest that metastases in other organs are immunologically “colder” than primary tumors^17,18^. The heavy involvement of immune cells led us to conduct more systematic investigation, and an unbiased clustering of all bone metastases using frequencies of various immune cells revealed three major clusters that are characterized by enrichment of macrophages/osteoclasts, monocytes, and regulatory/exhausting T cells (**Figure 1D** and **Figure S1G**). Indeed, the percentage of myeloid cells and the percentage of T cells exhibited a strong negative correlation across all cancer types (**Figure 1E**), which is not observed in normal control samples (**Figure S1H**). ^20–22^Both myeloid cells and T regulatory cells are immunosuppressive and support evasion of immunosurveillance^23^. However, their inverse correlation suggests parallel mechanisms selected by different tumors during their evolution to survive selective pressure exerted by the immune microenvironment in bone. Furthermore, the absence of osteoclasts in bone metastases also suggests alternative bone remodeling mechanisms. Overall, these data indicate previously uncharacterized archetypes of metastatic ecosystems that employ distinctive strategies to cope with challenges during bone colonization. We hereby refer to these subtypes by their characteristic contents, namely “Mφ-OC” (enrichment of macrophages and osteoclasts), “Mono” (enrichment of monocytes), and Treg-Tex (enrichment of regulatory and exhausting T cells), respectively.

Remarkably, we observed a complicated scenario when aligning different tumor types to the three ecosystem archetypes (**Figure 1F**). Whereas breast cancer bone metastases are classified as either Mφ-OC or Mono, kidney and lung cancer bone metastases exhibit all three archetypes. Specifically, most kidney cancer bone metastases belong to Treg-Tex that lacks OC, but lung cancers evenly distribute across different archetypes. These observations suggest both convergent and divergent evolution pathways – cancers stemmed from different tissues-of-origin can evolve similar colonization mechanisms, and conversely, cancers the same type also develop parallel mechanisms to colonize the same organ, bone.

### Myeloid and T lymphocytes have distinct developmental routes in different archetypes

Since immune archetypes are delineated by myeloid cells and T lymphocytes, we hypothesized that macrophages, osteoclasts, regulatory CD4+ T cells, and exhausted CD8+ T cells may exhibit unique developmental dynamics within these archetypes. Utilizing the *Dynamo* package for RNA velocity analysis^24^, we adopted the embedded least action path (LAP) principle to ascertain optimal lineage transition routes, shedding light on the formation kinetics of the immune archetypes. In the Mφ-OC archetype, our analysis revealed monocytes and macrophages as primary precursors for osteoclastogenesis. In contrast, monocytes in other archetypes display limited differentiation potential toward osteoclasts (**Figure 2A**). While gene expression velocities are consistent across archetypes (**Figure S2A**), myeloid-specific gene acceleration patterns in the Mφ-OC archetype show a progressive increase in gene expression turnover (**Figure S2B**). Moreover, myeloid cells in this archetype undergo a complex series of seven transcription factor (TF) activation phases along the LAP, indicative of extensive reprogramming and evolutionary TF profile shifts (**Figure 2B**).

**Figure 2.**
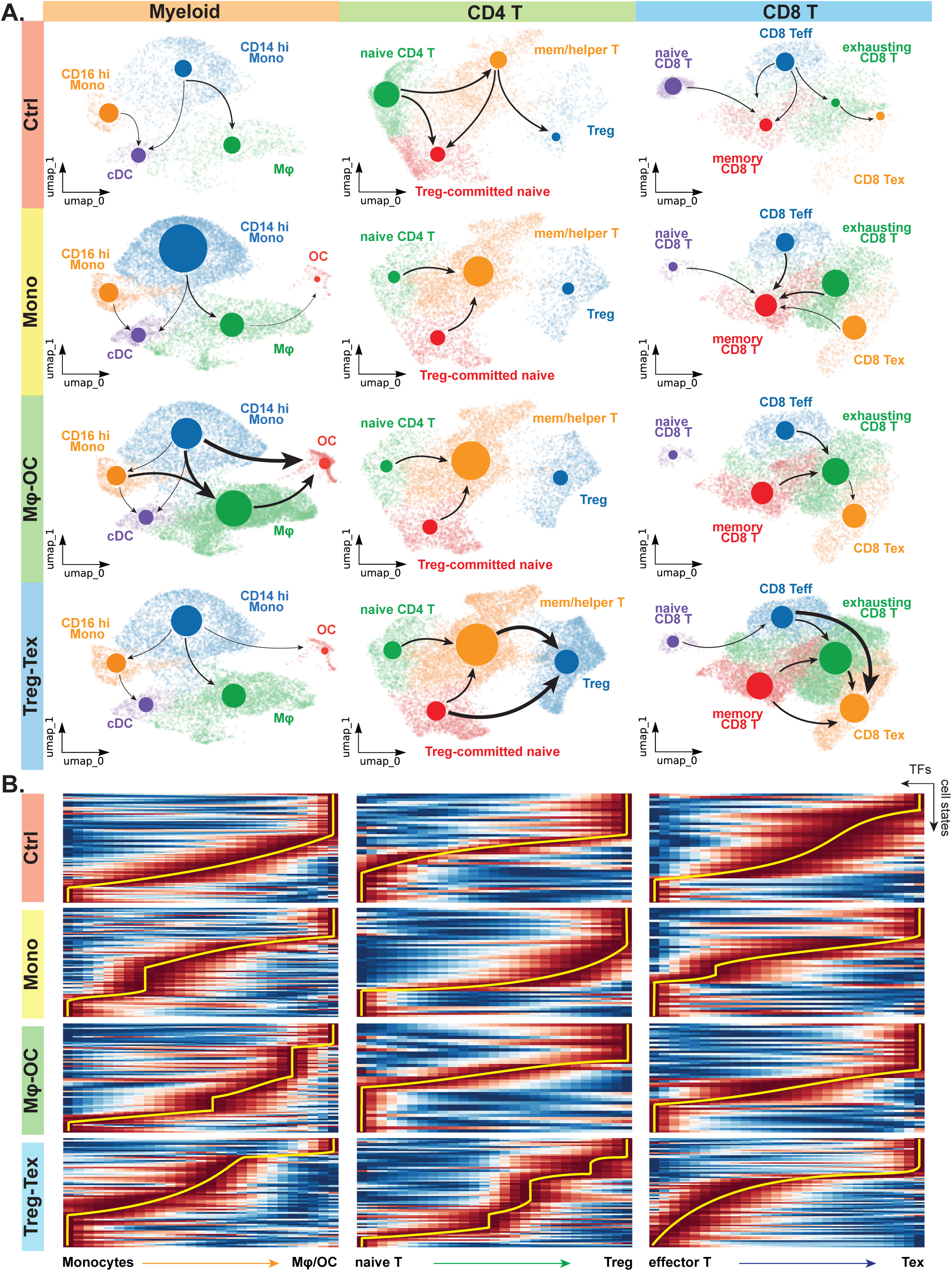
Myeloid and T lymphocytes cell fate characterization. **A.** Inferred trajectory analysis for myeloid cells, CD4, and CD8 T lymphocytes reveals distinct lineage commitment across different immune archetypes. Columns represent the three cell lineages; rows represent immune archetypes and control group. Arrows represents the cell state transition in a vector field projected on UMAP. Arrow thickness represents the transition probabilities, dot size denotes the cell population. Figures were produced from modified state graphs from Dynamo, projected on UMAP. **B.** Kinetic heatmap showing the human transcription factor (TF) expression kinetics across cell transitions from developmental initial to terminal states (restricted to those TFs that were used for cell state transition estimation). Distinct TF expression patterns are present in each immune archetype. TF waves are highlighted by yellow lines.

For CD4+ T lymphocytes, a subset of naïve T cells are predisposed toward a regulatory T cell fate, fueling Treg expansion in the Treg-Tex archetype—termed “Treg-committed naïve” cells (**Figure 2A**). These cells demonstrate distinct gene acceleration patterns, which appear to be interrupted in Mono and Mφ-OC archetypes but relatively more continuous and complete in Treg-Tex (**Figure S2B**). Their TF kinetics suggest substantial reprogramming within CD4+ T lymphocytes. On the other hand, CD8+ T lymphocyte analysis suggests a rapid transition from cytotoxic effector cells (Teff) toward an exhaustion state, even skipping some of the intermediate states (**Figure 2A**).

These data indicate that TF programs unique to each archetype govern the divergent developmental trajectories of the immune system, playing a crucial role in the establishment of each archetype. Nonetheless, these immune compartments interact within the metastatic microenvironment, influenced by both the cancer cells and the bone stroma. The implications of these interactions will be explored in the subsequent sections.

### Genomic copy number alterations in cancer cells are associated with ecosystem archetypes

To distinguish epithelial cancer cells from normal cells, we used the *infercnv*^25^ method to deduce copy number alterations (CNA) using single-cell transcriptome (**Figure 3A**). This approach successfully identified cancer cells as expected. Furthermore, it illuminated a wide range of CNA across individual bone metastasis (**Figure 3B** and **S3A**). Across different cancer types, high CNA is more frequently observed in breast cancers and the Mφ-OC archetype (**Figure 3C**). Further analyses revealed a positive correlation between CNA and fraction of macrophages/osetoclasts and an inverse correlation between CNA and fraction of monocytes particularly in breast cancer (**Figure 3D**), suggesting that monocyte-to-macrophage differentiation might be enhanced in breast cancer with increased genomic instability.

**Figure 3.**
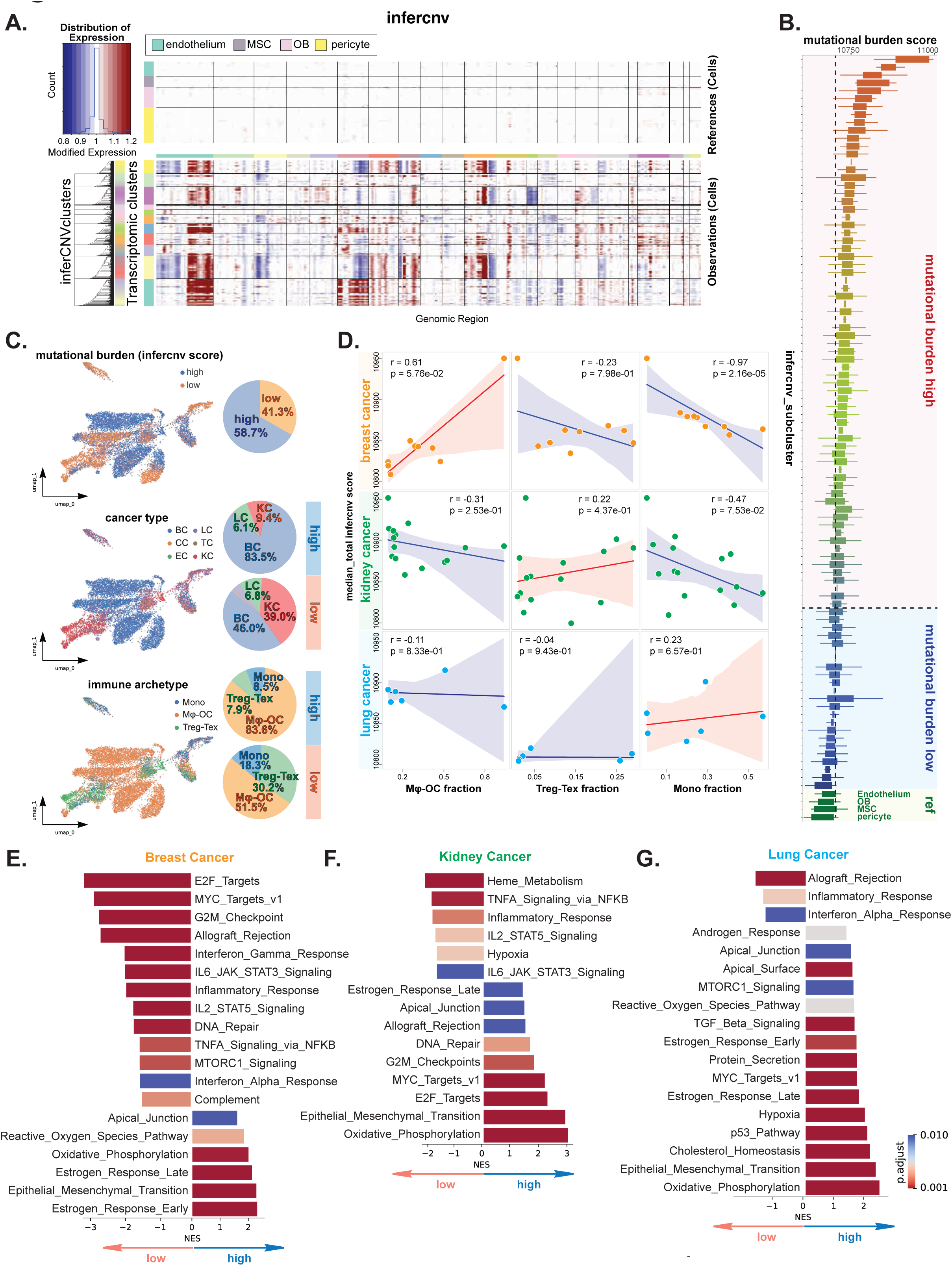
Characterization of epithelial/cancer cells and their correlation with immune archetypes. **A.** Epithelium copy number variation (CNV) estimation. Endothelium, MSC, OB, and pericytes were selected as references in epithelium copy number variation estimations. **B.** Assessment of Mutational Load. For each cell, mutational burden was quantified based on CNVs of genes. A scoring system was implemented whereby an increase of two or more copy numbers was allocated two points, a loss of two copy numbers received two points, and a single copy number change—either gain or loss—was assigned one point. Genes with no CNV were not scored. The cumulative scores for cells within each infercnv cluster were computed and then displayed in a bar chart, ordered from highest to lowest. Clusters were designated as having high or low mutational burden relative to a benchmark: the median score of a cluster exceeding the 75th percentile of scores in the reference cell population was considered to have a high mutational load. **C.** Distribution of CNV scores across cancer types and immune archetypes. The CNV mutational burden in epithelial cells is effectively illustrated using a UMAP. Pie charts on the right side highlight the prevalence of epithelial cells with either high or low mutational burdens across various cancer types (middle) and immune archetypes (lower). **D.** Correlation Matrix of CNV Scores and Cell Frequencies. This matrix plot illustrates the relationships between CNV scores and the frequencies of key cell types (macrophages, osteoclasts, regulatory and exhausted T cells) across different patients, organized by cancer type (rows) and immune archetype (columns). All correlations were analyzed using Spearman’s method. **E. F. and G.** Gene set enrichment analysis (GSEA) for the differentially expressed genes between the mutational high and low epithelium. This analysis was performed for breast, kidney, and lung cancers, identifying pathways with significant differences (adjusted p<0.01, by BH method).

Unbiased comparison within each cancer type revealed a common enrichment of inflammation-related pathways (e.g., STAT3/5, interferon pathways) in low CNA metastases across breast, kidney and lung cancers (**Figure 3E-3G**), suggesting more active immune responses against these tumors. On the other hand, the lack of these pathways in high CNA metastases suggests stronger immunosuppression mechanisms. Notably, a common pathway enriched in high CNA metastases of all three cancer types is oxidative phosphorylation (**Figure 3E-3G**), which implies linkage between metabolic changes and genomic instability in bone metastases.

In search for cancer cell-intrinsic determinants of immune archetypes, we compared transcriptomic profiles of cancer cells. Interestingly, we observed that estrogen responses appear to be elevated in the Mφ-OC archetype in both breast and kidney cancers (**Figure S3B-S3F**). It should be noted that all 9 breast cancers included in this study were derived from ER+ primary diseases. The sustained ER signaling in bone metastases may result from evasion mechanisms evolved in response to anti-estrogen therapies and appear to be associated with a Mφ-OC-enriched as opposed to monocyte-enriched environment. For kidney cancer metastases, the estrogen response is unexpected and may be related to estrogen activities in macrophages, which will be elaborated in a later section. Reciprocally, pathways that are commonly depleted from Mφ-OC and enriched in other archetypes include G2M checkpoint, Myc targets and E2F targets, which collectively suggest more active cell cycling. Overall, these analyses support that the ecosystem archetypes may result from cancer cell-intrinsic characteristics that co-evolve with the bone microenvironment.

### Other cells in different archetypes

Having examined immune and cancer cells, we asked if other cell types also exhibit significant differences among different archetypes. Our SVM-based annotation successfully identified endothelial cells, mesenchymal stem cells (MSCs), osteoblasts (OBs), pericytes, and fibroblasts (**Figure 4A**), all of which have been implicated in metastatic colonization^26^. Like immune cells, we also manually validated the annotation using widely used marker genes (**Figure 4B**).

**Figure 4.**
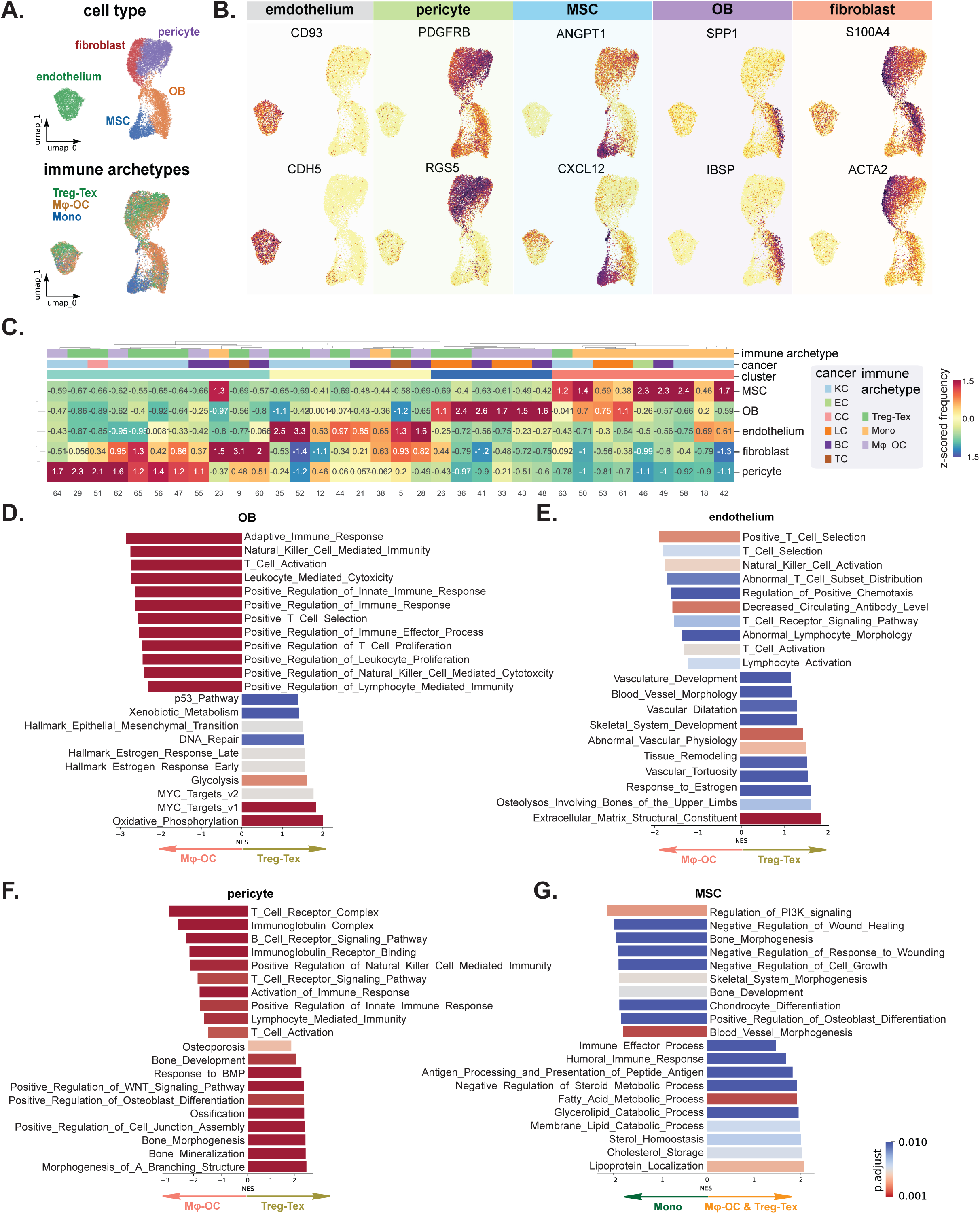
Bone stromal niche characterizations. **A.** Major cell types and stromal cell immune archetype identities in UMAP. **B.** Validation of major cell-specific marker expression in stromal cell populations. **C.** Supervised stromal cell clustering based on z-scored cell frequencies in each patient. Patients’ archetype and cancer types were annotated. **D. - G.** Gene set enrichment analysis (GSEA) between the immune archetypes among all stromal cells, with unbiased pathway selection based on a stringent cutoff (adjusted p<0.01, by BH method).

We first analyzed if frequencies of these stromal cells vary across the three immune archetypes (**Figure 4C**). Indeed, the most significant finding with this regard is the abundance of MSCs in the Mono archetype. The association of other cell types with immune archetypes is much less notable, suggesting similar constitution of non-immune, non-cancer compartment.

We next characterized the phenotypic disparity between the same cell types in different immune archetypes. We focused on differences between Mφ-OC and Treg-Tex because of the negative correlation between macrophage/osteoclasts and regulatory/exhausted T cells (**Figure 4D-4F**). The comparison of MSCs is an exception because of their predominant association with the Mono archetype. We therefore did the analysis between Mono vs. the other two archetypes (**Figure 4G**). A number of interesting findings were made. First, osteoblasts in Treg-Tex archetype appear to be regulatory to immune responses, but they exhibit a number of other functions such as estrogen responses (**Figure 4D**) in Mφ-OC. Second, similar to osteoblasts in Treg-Tex, endothelial cells in this archetype also seems to play roles in immune regulation, which is in contrast to endothelial cells in the Mφ-OC archetype. The latter enriches vascular modeling and development pathways along with a pathway denoting “response to estrogen”.

Third, pericytes in Treg-Tex, again, exhibit immunoregulatory phenotype. In contrast, their counterparts in Mφ-OC appear to be promoting osteoblast differentiation and ossification (**Figure 4F**), which is consistent with our previous conclusions drawn from Mφ-OC models. Finally, MSCs in the Mono archetype appear to orchestrate lipid metabolism as opposed to pathways associated with indicate chondrocyte and osteoblast differentiation in other archetypes, which indicates that these multi-potent stem cells may be skewed toward divergent directions of differentiation (**Figure 4G**).

Taken together, these results support that the archetypes defined by their distinctive immune landscape also differ in other cell compartments. Various cell types seem to constitute networks involving intricate crosstalk as tumors progress. The interactive co-evolution in turn results in divergent functions of each cell type. Thus, the roles of a specific cell type cannot be precisely understood without insights into the entire metastasis ecosystem.

### The in-depth characterization of immune cells identifies ER signaling as a determinant of macrophages associated with osteoclast-driven metastases

Having analyzed the divergence between different archetypes, we next examined the microenvironmental diversity within the same archetypes. Principle Component Analysis (PCA) was performed on scRNA-seq profiles of individual patients and healthy control samples. Cells of the same type were then combined and subjected to PCA across patients in each immune archetype (**Figure 5**). In this analysis, the distance between any two points or the area of a polygon connecting multiple points indicate the degree of dissimilarity among the underlying data. Therefore, when all points of the same cell type were connected, the covered area reflects patient-to-patient variations of this cell type, which were also validated and ranked based on sum of eigenvalues on both PC1 and PC2 (**Figure S4A**). Compared to their normal counterparts (**Figure 5A**), microenvironmental components of bone metastases exhibited dramatic variations in all three archetypes (**Figure 5B-4D**). Some of the variations even exceed those of epithelial cells (cancer cells). This observation indicates that metastatic seeds of different tumors may sculpt the bone marrow soil and induce significant phenotypic plasticity even within the same archetype. Of note, our companion study also made a similar observation and experimentally tackled the underlying mechanism.

**Figure 5:**
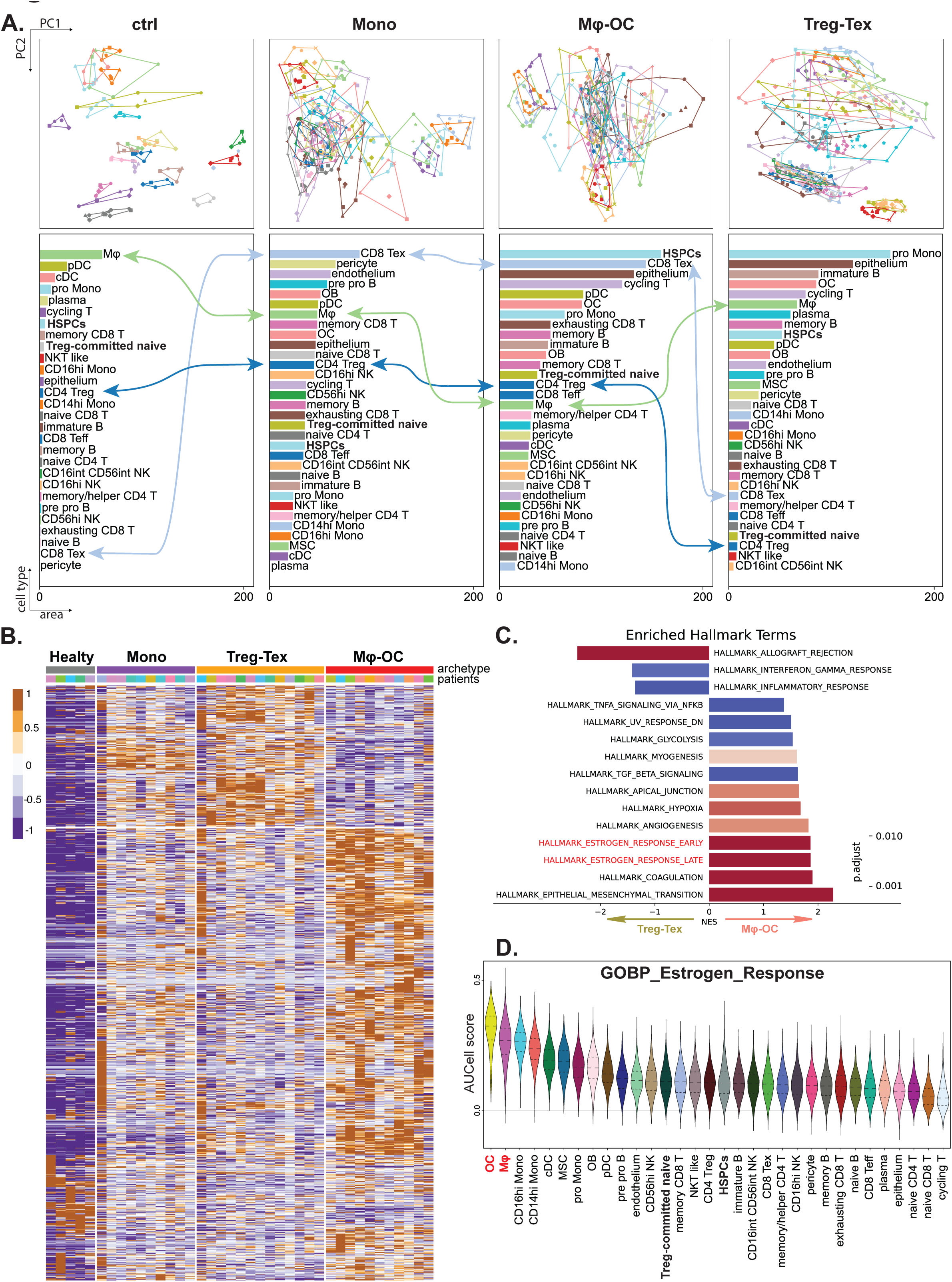
Comparative transcriptomic variance analysis and macrophage profiling between immune archetypes. **A.** PCA of aggregated transcriptomic data (pseudo-bulk) by cell type. In these plots, each data point, colored to represent different cell types, corresponds to patients from various cancer types or healthy control donors. Lines connecting the points encircle regions on the plot, with the size of these areas indicating the degree of inter-patient variance within each cell type. The area from each cell type is ranked in descending order in each bar plot, showing the transcriptomic heterogeneity changes between different archetypes. **B.** Heatmap showing significant differentially expressed genes (DEGs, log2 fold change > 1.5, p-value < 0.05) by comparing macrophages in the Mφ-OC group versus those in Treg-Tex patients. **C.** GSEA results from the pathway analysis. All genes that were differentially expressed in macrophages across Mφ-OC and Treg-Tex were applied for enrichment analysis. Reference pathways are from the “Hallmark” category in the MSigDB database. P-values adjusted by BH method. **D.** Pathway “GOBP_Estrogen_Response” activity was accessed in every cell by calculating scores based on expression levels of all pathway-specific genes. Wilcoxon rank-sum test was used. Cells were ranked by the medium score.

Further investigation revealed that the defining components of each archetype typically exhibit lower diversity in this archetype compared to others. For instance, CD8ex cells appear to be highly heterogeneous in Mono and Mφ-OC, but the heterogeneity is much reduced in Treg-Tex metastases. And the same is true for Mφ and monocytes in Mφ-OC Mono, respectively (see lines connecting these cell types between different categories in Figure **5A-5D**). These observations support that the dominant cell types of each ecosystem evolve conserved or convergent functions, which probably represent the driving force of bone colonization.

Subsequently, we analyzed macrophage-specific differential gene expression profiles across the archetypes (**Figure 5B**), revealing distinct transcriptomic patterns. The phenotypic diversity of macrophages has been intensively studied in previous studies ^27,28^. Here, our data provide insights into the how such diversity is inter-related with entire metastasis ecosystem.

To further dissect the functional and pathway distinctions, we conducted Gene Set Enrichment Analysis (GSEA) on differentially expressed genes (DEGs) (**Figure 5C**). Remarkably, estrogen-related pathways were significantly enriched in the Mφ-OC archetypes. Additionally, osteoclasts and macrophages exhibited the highest scores for estrogen-responsive pathways among all cell types (**Figure 5D**), regardless of the cancer types (**Figure S4A-S4C**) or gender (**Figure S4B**). This finding correlates with the enrichment of estrogen-responsive pathways in the epithelium of kidney cancer within the Mφ-OC archetype (**Figure S3C-S3D**). Collectively, these results underscore the importance of estrogen signaling in macrophages from Mφ-OC patients, across cancer types, potentially linked to macrophage-associated, osteoclast-driven metastases.

### Crosstalk between different cell types leads to divergent bone colonization and immuno-evasion mechanisms

We conducted cell-cell communication analyses to delineate mechanisms behind the evolution of different archetypes. Toward this end, we focused on the comparison between Mφ-OC and Treg-Tex archetypes as they appear to represent two opposite types of systems. Using the algorithm, *CellChat*, we estimated strengths of intercellular signaling in each cell type either as receiver (expressing receptors) or sender (expressing ligands) (**Figure 6A**). Interestingly, in both Mφ-OC and Treg-Tex archetypes, mesenchymal cell types, including pericytes, OB and MSCs, appear to be the strongest signal sources, whereas CD8 T cells are among the most prominent signal receivers (**Figure 6A**). This is consistent with many previous studies supporting the roles of MSCs ^29^. We subsequently quantified the relative probability of each signaling pathway across cell pairs within the Mφ-OC and Treg-Tex archetypes (**Figure S5A, Table S3**). We identified the key signaling pathways in **Figure 6B** that exhibit more than a 75% difference between the immune archetypes, with osteoclasts serving as the primary receptor cells. To validate the roles of signaling molecules identified in the Mφ-OC archetype, we performed a series of *in vitro* assays to evaluate their effects on osteoclastogenesis. CD14+ monocytes were isolated from healthy human peripheral blood and used as precursors for osteoclast induction. We introduced RANKL and M-CSF to initiate osteoclastogenesis and supplemented the culture with three selected signaling factors. The impact of these factors was assessed by measuring the expression of osteoclast signature genes, including TRAP (tartrate-resistant acid phosphatase), CTSK (cathepsin K), DC-STAMP (dendritic cell-specific transmembrane protein), and NFATC1 (nuclear factor of activated T-cells c1), using qPCR. Additionally, TRAP assays were conducted to visualize osteoclast formation. Our evaluations confirmed the roles of TWEAK, COMP, and NRP1 in enhancing osteoclastogenesis, likely through mechanisms mediated by macrophages.

**Figure 6.**
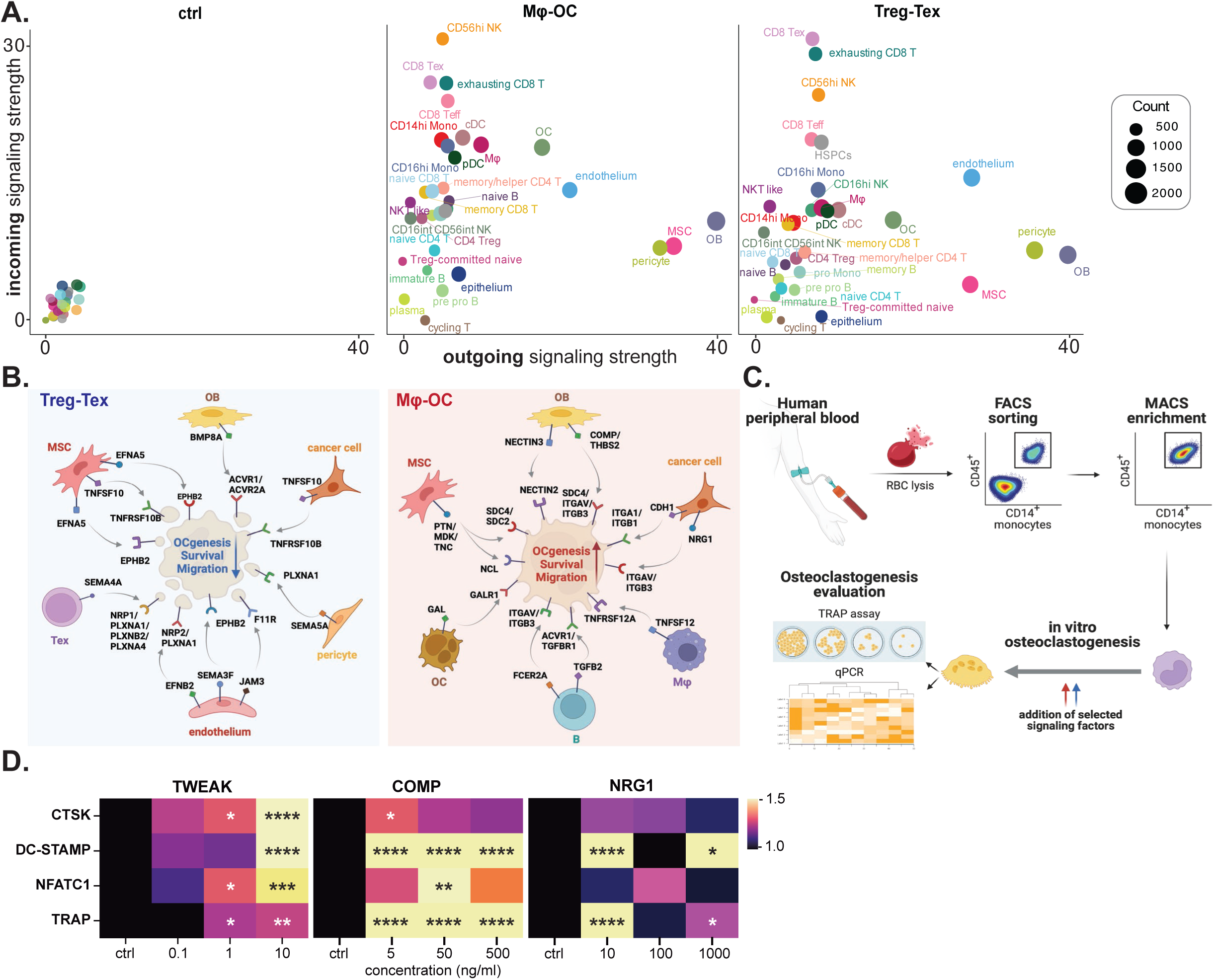
Analysis of cell-cell communication unveiling regulatory mechanisms in the formation of the Mφ-OC archetype. **A.** Analysis of cell-cell communication signal flow. Outgoing signal strength is shown on the x-axis and incoming signal strength on the y-axis, comparing the Mφ-OC and Treg-Tex archetypes with healthy samples serving as references. **B.** Identification of key ligand-receptor pairs. This analysis compares the relative signaling strengths between the Mφ-OC and Treg-Tex archetypes, focusing on osteoclasts as the signal receivers (from Figure S5A). In the Mφ-OC archetype, signaling pathways tend to promote and maintain osteoclastogenesis, whereas in the Treg-Tex archetype, the effects are generally inhibitory. **C.** *In vitro* experimental validation. Using human peripheral blood, CD14+ monocytes were isolated and cultured. Osteoclastogenesis was induced by adding RANKL and MCSF to the culture. Selected recombinant factors identified in part B were also incorporated to assess their impact on osteoclast formation. The efficacy of these interventions was evaluated through TRAP assays and qPCR, targeting osteoclast-specific gene expression. **D.** qPCR analysis of osteoclast signature genes. The expression levels of osteoclast-related genes CTSK, DC-STEMP, NFATC1, and TRAP were quantified using qPCR to evaluate the effects of selected signaling factors from the Mφ-OC archetype. The relative differential expression of these genes were calculated based on reference gene GAPDH, and presented as heatmap. In the Mφ-OC archetype, factors tend to enhance osteoclastogenesis, whereas factors from the Treg-Tex archetype show inhibitory effects. The experimental setup included control monocytes (ctrl) subjected to osteoclastogenic conditions. Treatments with each protein were tested at three concentrations in the culture medium to determine dose-dependent effects. Statistical significance was determined using one-way ANOVA, with significance levels denoted as follows: *p<0.05; **p<0.01; ***p<0.001; ****p<0.0001.

### The immune archetypes may underpin the distinct clinical behavior of bone metastases from different cancer types

We examined records of several cancer types treated at MD Anderson Cancer Center since the late 1990s (the beginning of bisphosphonates usage) and compared proportion of patients that required orthopedic surgeries for treatment of their bone metastasis, a hallmark of severe skeletal-related events (SREs). Strikingly, kidney cancer bone metastases exhibit a 5-7 fold higher risk of surgery (**Figure 7A-7C**). Admittedly, many factors vary between different cancer types, limiting their comparability. For instance, the bone metastasis rate and the survival time of bone metastasis patients are vastly different between breast, kidney, and lung cancers. Prior treatments other than bisphosphonates or denosumab may also alter disease progression in bone. Nevertheless, these factors cannot explain the disproportionally high incidence of severe SREs in kidney cancers. Using the comparison between breast and kidney cancers as an example: breast cancers have a higher bone metastasis rate (DOI: 10.3390/cells10112944) and longer survival time^30^, hence more patients living longer with bone metastases would benefit from orthopedic surgeries once SRE develop. However, only 6/1000 breast cancer patients need orthopedic surgeries as compared to 37/1000 for kidney cancer (**Figure 7C**). This discrepancy may be explained by the different constitution of bone metastasis ecosystems: kidney cancer bone metastases are predominantly the Treg-Tex or Mono archetypes, whereas breast cancers are mainly Mφ-OC archetype.

**Figure 7.**
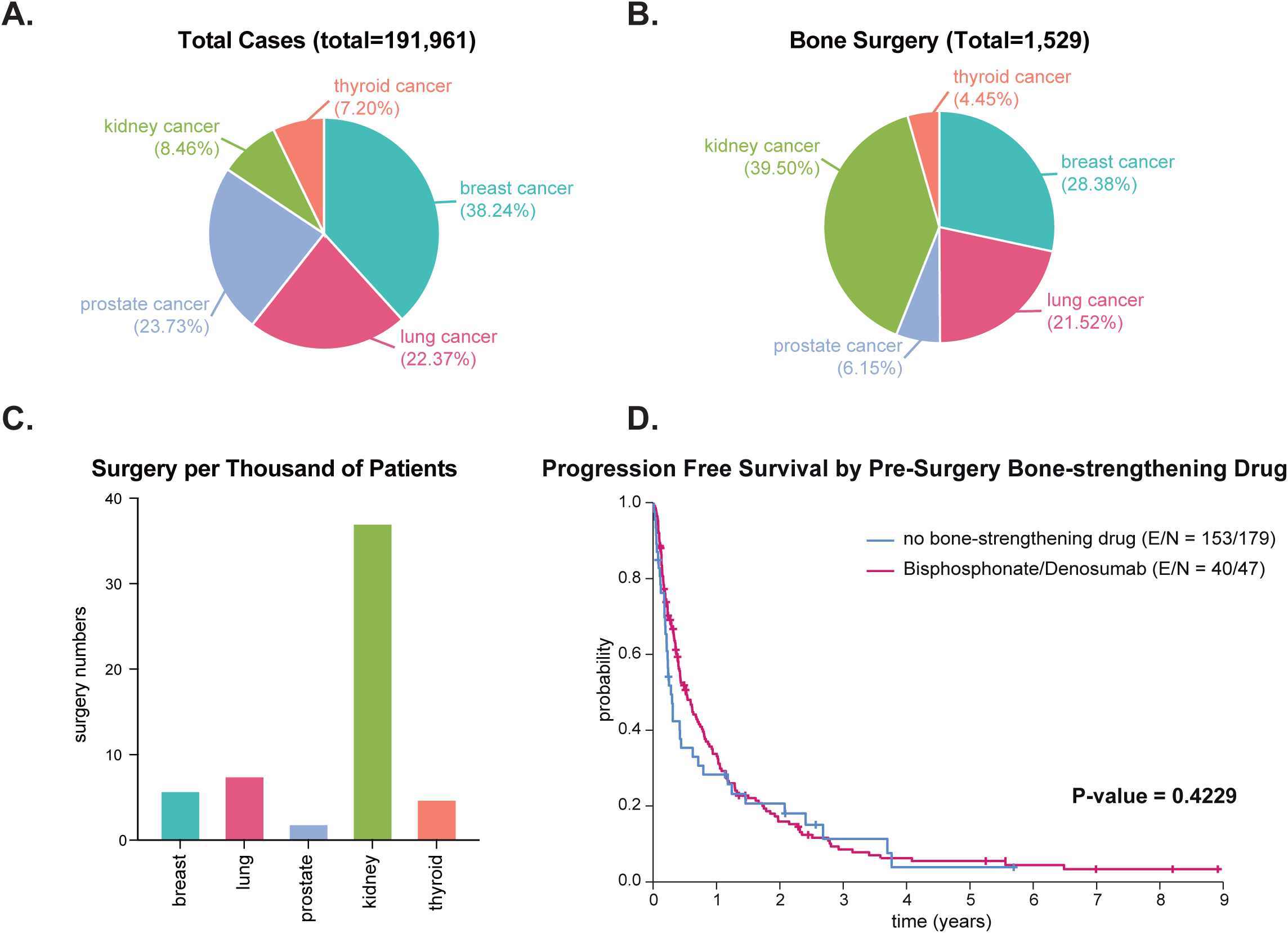
Distinct clinical behaviors of bone metastases from different cancer types. **A.** The case numbers of indicated cancer types diagnosed between 1998 and 2018 at MD Anderson Cancer Center **B.** The case numbers of indicated cancer types undergone orthopedic surgeries between 1998 and 2018 at MD Anderson Cancer Center. **C.** Numbers of orthopedic surgeries per 1000 patients of the indicated cancer types. **D.** Kaplan-Meier curves of progression free survival of kidney cancers with or without treatment of bisphosphonates and/or denosumab. The P values were calculated by the log-rank test.

For many cancer types including breast, agents targeting osteoclasts are often used to mitigate symptoms of bone metastases and prevent SREs. Indeed, numerous studies have demonstrated that bisphosphonates and denosumab both reduce SREs and prolong progression-free survival in breast cancer patients^31^. The efficacy of these agents, however, is insignificant for kidney cancer bone metastases in our database (**Figure 7D**), arguing against a major role of osteoclasts in these patients.

Taken together, these data highlight the need for better understanding of different bone colonization mechanisms undertaken by individual tumors – both between different cancer types and potentially also among patients of the same cancer type. Data shown in this study represent the first systematic endeavor toward this direction and lay foundation for future mechanistic and therapeutic research.

## DISCUSSION

The mechanism by which different cancer types colonize the same organ has not yet been systematically investigated. Our data revealed three archetypes of ecosystems characterized by different immune landscapes. The development of these ecosystems represents divergent evolution propelled by two major sources of selective pressure – immunosurveillance and confinement of mineralized bone tissues. For the former, different ecosystems develop parallel immunosuppressive mechanisms primarily based on T regulatory/exhausted cells, ER+ macrophages, and monocytes, respectively. For the latter, different ecosystems diverge on the driving force of metastasis-induced bone remodeling. Activated osteoclasts are enriched in the Mφ-OC type and potentially also the Mono type, which encompass almost all breast cancers.

This is consistent with the large body of literature demonstrating the essential roles of osteoclasts in osteolytic bone metastases. However, the Treg/Tex type lacks osteoclasts and is unlikely to rely on osteoclasts for bone remodeling. Kidney cancer is the major cancer type in this class. In fact, evidence has emerged that kidney cancers invade bone matrix by inducing apoptosis of osteocytes rather than activating osteoclasts^32^. In the clinic, osteoclast-targeting agents, such as bisphosphonates, are not as effective on kidney cancer bone metastasis as they are in other cancer types^16^, consistent with the fact that osteoclasts are not the major driver of bone colonization.

Interestingly, besides the divergence across different cancer types, we also observed that individual bone metastases of the same cancer type evolve vastly different mechanisms. For example, the six lung cancer bone metastases examined in our study evenly distributed across all three types of ecosystems. The determinants that sculpt metastatic evolution toward one ecosystem over the others remain to be investigated. One factor examined in our study is the amount of CNAs (**Figure 2**): the Mac/OC ecosystem is associated with high CNA, and this association can explain the diversity of breast cancer bone metastases. Further studies will be needed to establish causal links between overall frequency of genetic variations and the archetype of bone metastasis ecosystem.

Conversely, different cancer types can also evolve into similar ecosystems and utilize the same bone colonization mechanism. This may not be surprising considering the aforementioned selective pressure in the BME that is commonly encountered by metastatic cancer cells.

However, it is curious to ask why cancers from different tissues of origin end up choosing the same mechanism among three parallel options. Again, if the level of genomic variation is a key determinant, then how is this variation sensed by the BME and how does it trigger the development of a particular type of ecosystem. It is possible that DNA damage response pathways and/or tumor-specific antigens are involved in the divergence among different mechanisms, which warrants further investigations.

Our data suggest that the mesenchymal stromal cells (including mesenchymal stem cells, fibroblasts and pericytes) are important sources of signals in all bone metastases, although the specific signals and the major recipient cell types may differ between different ecosystems as revealed by the cell-cell communication analyses. These disparate mesenchymal signals together with the vastly varying immune milieu appear to converge on opposite regulation of osteoclast activation: whereas the fibroblast-derived periostin, the MSC-derived CSF1 and CD4fh-derived TWEAK promote osteoclastogenesis in the Mφ-OC ecosystem, the exhausted T cells may inhibit the same process in the Treg-Tex ecosystem. Thus, the complicated, multi-cell type crosstalk seems to dictate whether or not the bone colonization is osteoclast-dependent.

Immune cell infiltration has been considered a hallmark of immunogenic tumors that tend to be responsive to immune checkpoint blockade (ICB). Recent comparisons between matched human primary and metastatic tumors, few of which were from bone, suggest that metastasis is typically “colder” and even less immunogenic^17,18,33^. Herein, our data revealed that bone metastases of many cancer types are in fact well infiltrated by immune cells. Despite their presence, however, these cytotoxic cells are blunted by different immunosuppressive mechanisms in different types of ecosystems.

## AUTHOR CONTRIBUTIONS

X.H.-F.Z. and Z.X. conceived the concept and designed the experiments. Z.X. conducted and analyzed the animal studies, flow cytometry, and imaging experiments. F.L. performed the single-cell RNA sequencing and bioinformatic analysis. Y.D., T.P., Z.G., and R.L.S. aided in the collection of clinical samples. Y.W. assisted in performing intra-iliac injections. X.H. and Y.G. assisted in performing intra-cardiac injections. J.L., I.L.B., and W.Z. contributed to maintaining mouse strains. W.L., L.Y., D.G.E., and S.A. provided assistance with animal work. H.L.C. aided in cell differentiation work. M.W.D., E.C., and other authors contributed to manuscript editing. X.H.-F.Z. supervised the research.

## SUPPLEMENTARY FIGURE LEGENDS

**Supplementary Figure 1.**
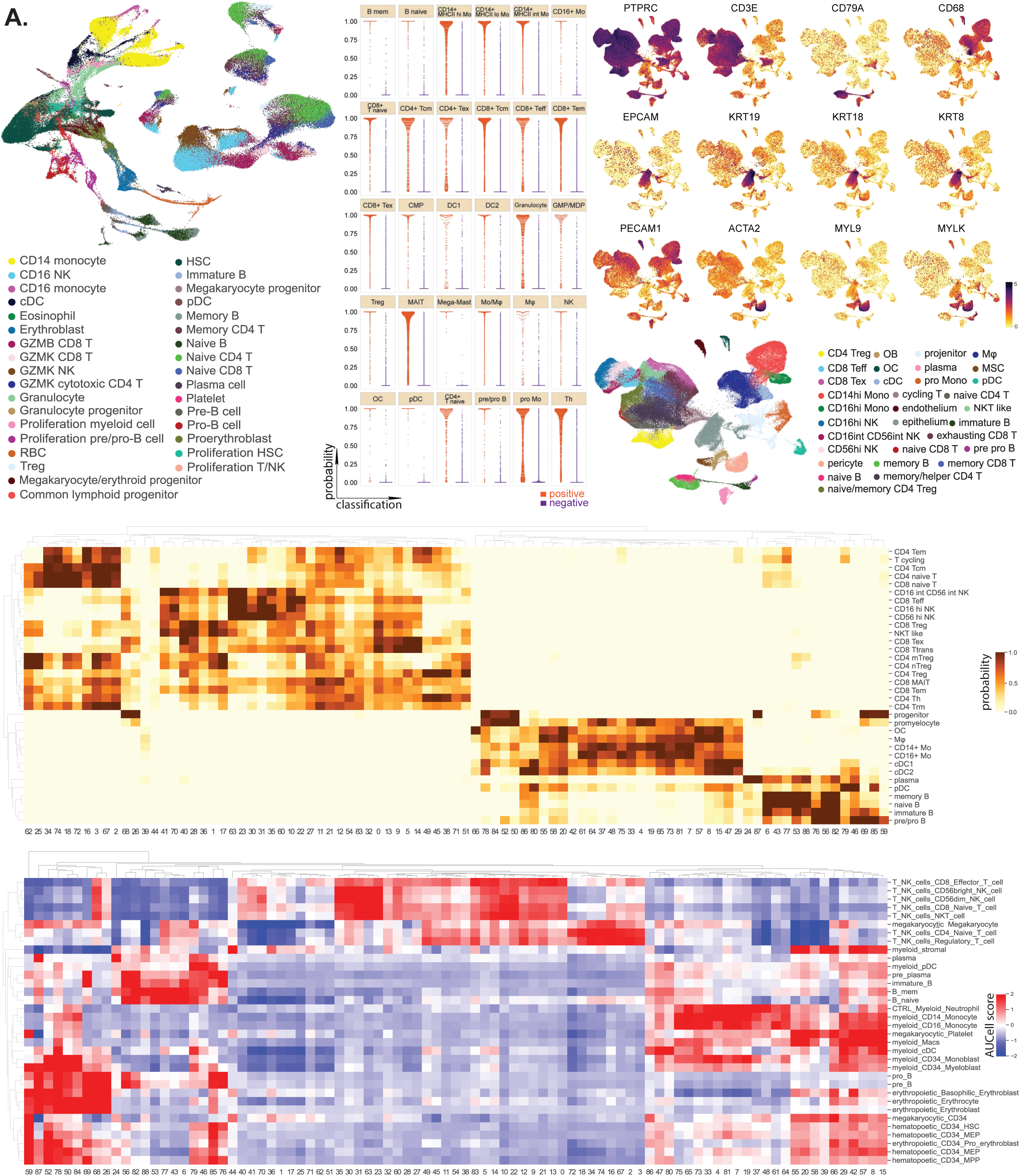
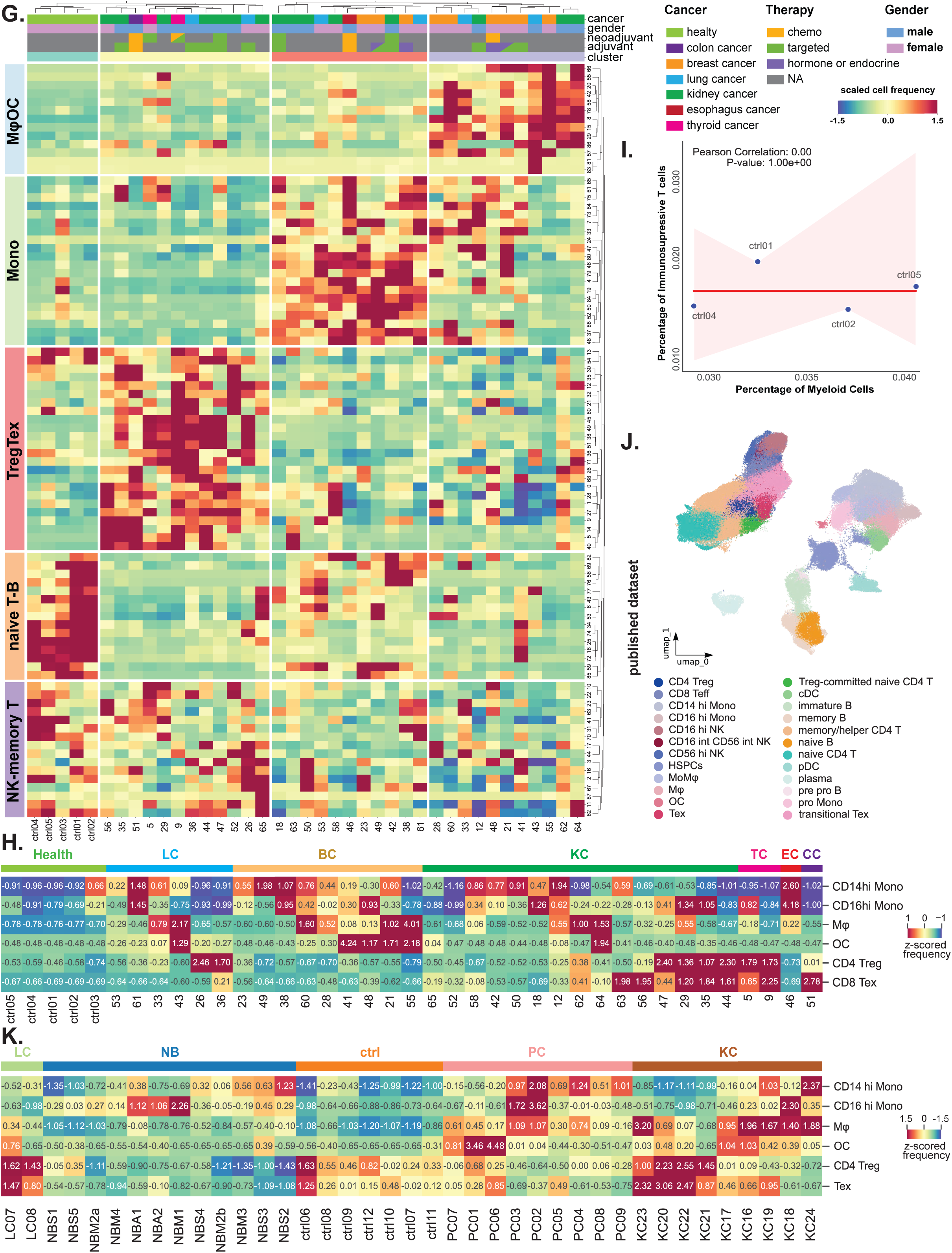
Cell type annotation using SVM trained classifier and annotation validation. As related to Figure 1. **A.** SVM training was performed using an integrated single-cell reference dataset to facilitate cell type annotation, as detailed in the Materials and Methods section. **B.** Scatter plot assessing the quality of the SVM classifier, trained on the reference dataset, showing classification performance. Positively identified cells are marked with red dots, while negatively identified cells are marked with purple dots. **C.** Heatmap displaying the classification probabilities for each reference cell type against dimension reduction clusters at resolution 5 in the metastatic cohort dataset, post-prediction. **D.** Feature plots illustrating the expression of classical marker genes across immune cells (first row), epithelial/cancer cells (second row), and stromal cells (third row). **E.** UMAP visualization depicting the SVM-predicted cell types within our bone metastasis patient cohort. **F.** Verification of SVM-predicted cell types using classical markers from flow cytometry, bulk sequencing data, and additional single-cell sequencing datasets. AUCell scores for specific gene sets were computed for each cluster at resolution 5, serving as cross-validation for the SVM predictions shown in panel C. **G.** Heatmap of unsupervised patient clustering based on scaled cell frequency data. This visualization uses data from clusters at dimension reduction resolution 5, with additional patient information such as gender and clinical therapy indicated in the heatmap header. **H.** Analysis of patient clustering by cell type frequencies. This analysis focuses on specific cell types including monocytes, macrophages, osteoclasts, and regulatory/exhausted T cells. Notably, among healthy control samples, no correlation was observed between the frequencies of myeloid cells and regulatory/exhausted T cells.

**Supplementary Figure 2.**
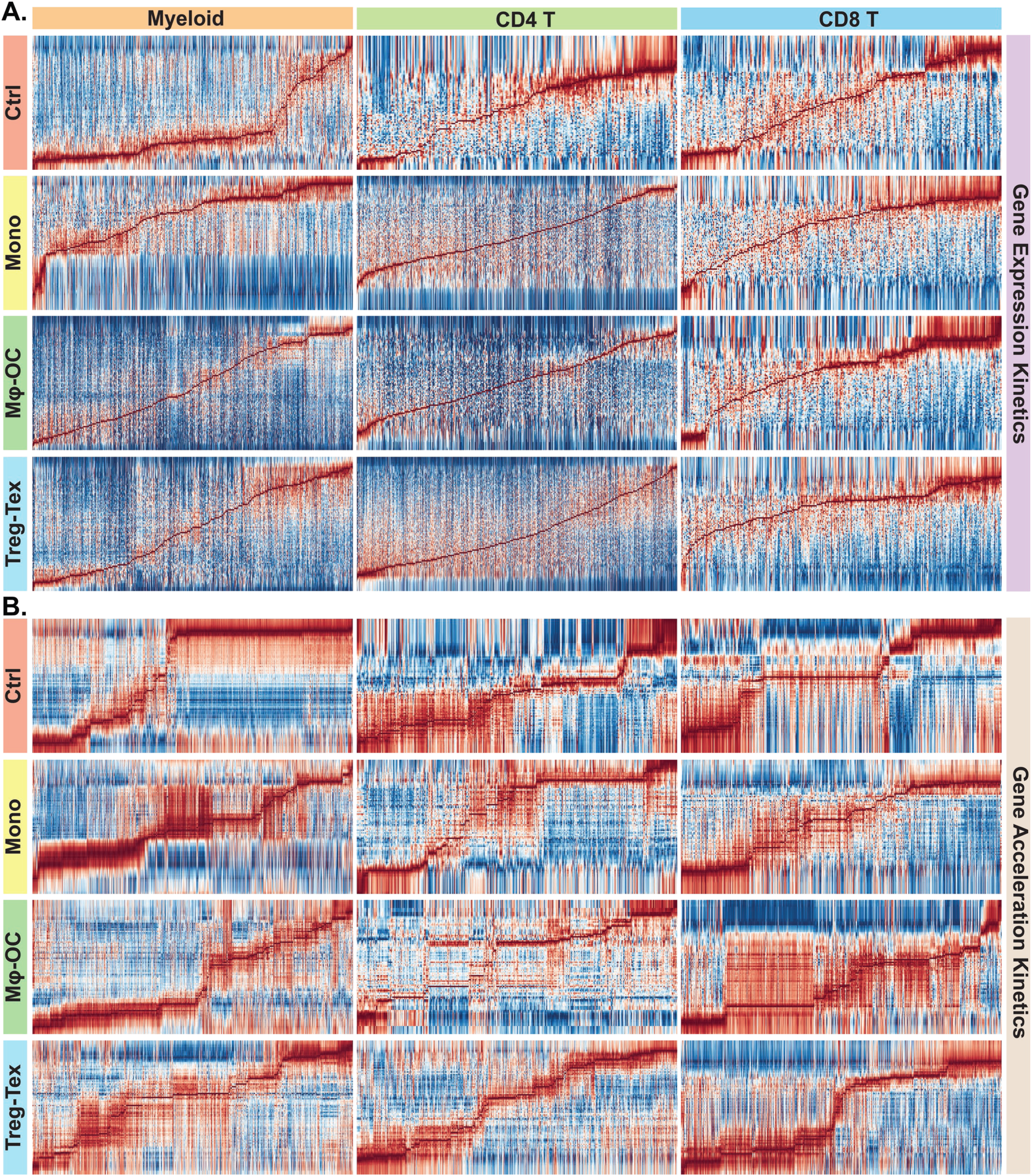
Gene expression and acceleration kinetics from RNA velocity analysis. As related to Figure 2. **A.** Gene expression kinetics: This series of heatmaps displays the expression velocity profiles of various genes (rows) across different cell states (columns) within distinct lineages and immune archetypes. **B.** Gene acceleration kinetics: These heatmaps illustrate the rates of gene transcript processing changes—including splicing and degradation—across different cell states (columns) in various lineages within different immune archetypes.

**Supplementary Figure 3.**
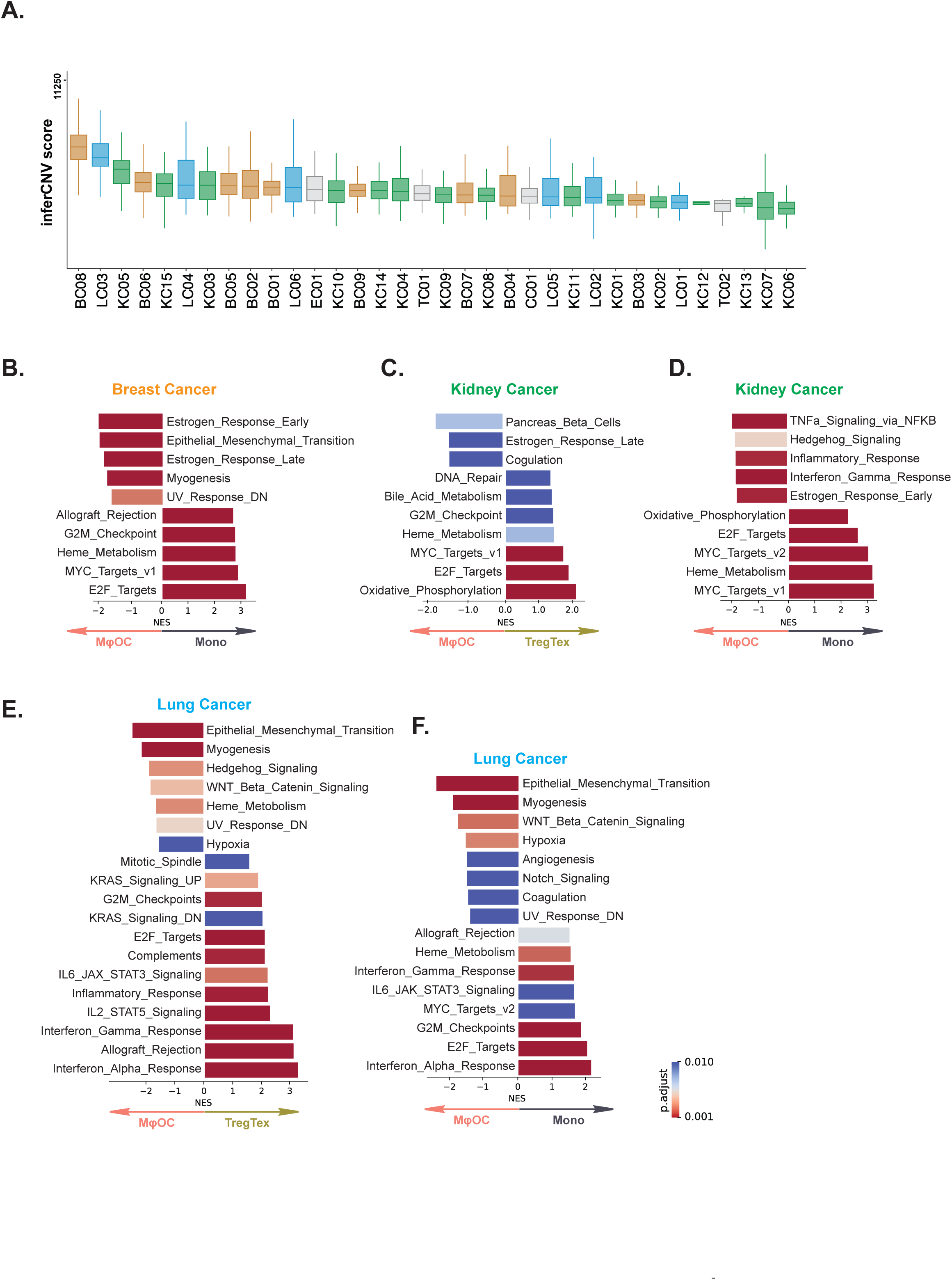
Epithelium mutational burden calculation and pathways analysis. As related to Figure 3. **A.** Per-Patient infercnv Mutational Scores. This plot ranks each patient based on the aggregated infercnv scores from all cells within that patient. Different cancer types are indicated by varying colors. **B. -F.** Gene Set Enrichment Analysis (GSEA) of GO Biological Processes. This series of analyses compares epithelial cell GOBP pathways across different immune archetypes within each cancer type. Pathways with significant differences (adjusted p<0.01, by BH method) were selected and displayed.

**Supplementary Figure 4.**
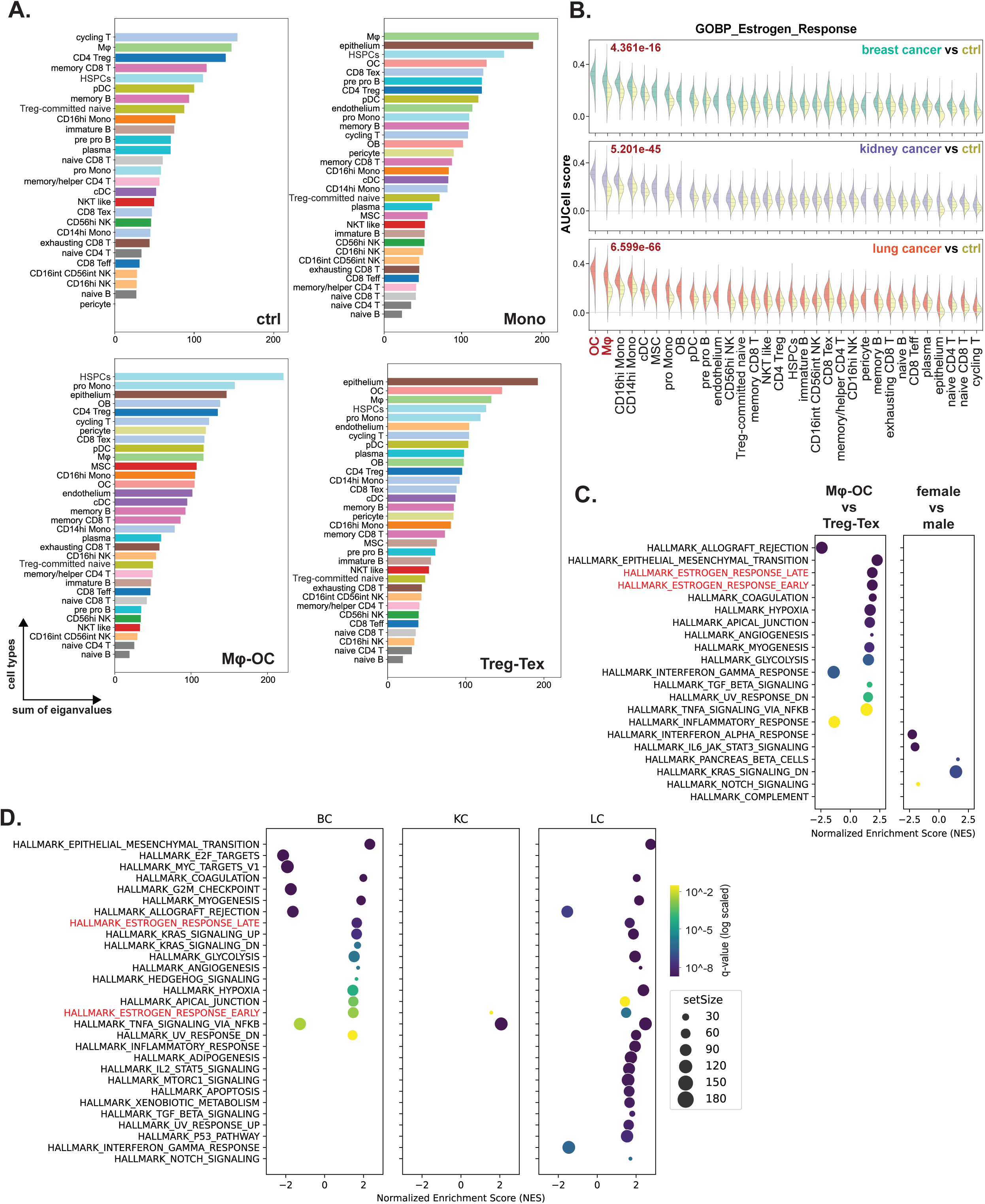
Additional transcriptomic variance calculation based on PCA and estrogen pathways activities in myeloid lineages. As related to Figure 5. **A.** Calculation of Transcriptomic Variance: This analysis involves summing the eigenvalues derived from PCA, which quantify data variance across PC1 and PC2, to represent the overall transcriptomic variance for each cell type across all patients. The summed eigenvalues for each cell type are then displayed in a bar chart in descending order to illustrate varied degree of inter-patient heterogeneity. **B.** We computed the pathway “GOBP_Estrogen_Response” activities using AUCell method and ranked the cell type by the medium scores. Furthermore, the violin plot was split into half as to compare different cancer types to healthy control groups. The p values (Wilcoxon Rank Sum Test) for the osteoclasts in cancer vs control samples were indicated. **C.** Cross-comparison of GSEA results within macrophage populations, delineated by immune archetypes (Mφ-OC vs. Treg-Tex), gender (female vs. male), or the cancer types with the largest patient cohorts (breast cancer vs. kidney cancer). The y-axis displays shared Hallmark pathway terms, with estrogen-related pathways highlighted in red when comparing immune archetypes or cancer types. **D.** Comparison within macrophage populations looks at patients from the three major cancer types, categorized by higher (patients belongs to Mφ-OC archetype) or lower (patients belongs to Treg-Tex archetype) macrophage fractions. The “Hallmark_Estrogen_Response_Early” pathway appears consistently across all three cancer types.

**Supplementary Figure 5.**
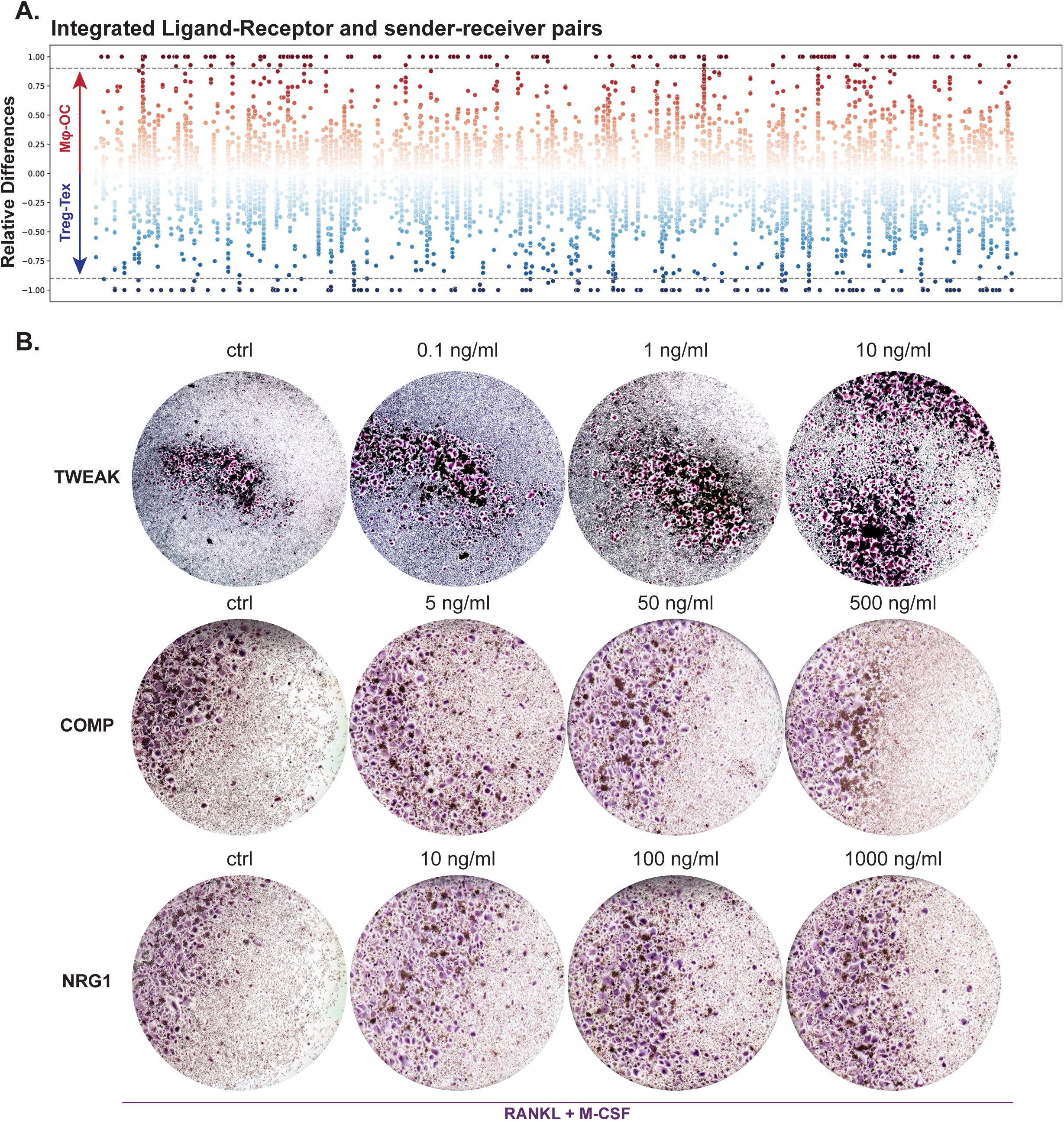
Validation of Signaling Molecule Effects on Osteoclastogenesis. As related to Figure 6. **A.** Comparative analysis of ligand-receptor activities. This scatter plot illustrates the relative differences in ligand-receptor pair activities between the Mφ-OC and Treg-Tex archetypes. Each column represents a distinct ligand-receptor pair, and each dot symbolizes a cell pair with a fixed direction of information flow (sender->receiver, e.g., Mφ->pericyte, where Mφ->pericyte is not equivalent to pericyte->Mφ). Positive values indicate that specific signaling pathways are enriched in cell pairs from the Mφ-OC archetype, while negative values are associated with the Treg-Tex archetype. Scaled relative differences are displayed numerically, with potential archetype-specific signaling pathways and cell pairs identified based on the top 90% of differences, marked by dotted lines. **B.** Validation of signaling factor effects on osteoclastogenesis. TRAP assay was employed to evaluate the impact of selected signaling factors on osteoclastogenesis. Purple-stating signaling within the assay wells indicates a positive response in osteoclast formation.

## STAR METHODS

### KEY RESOURCES TABLE

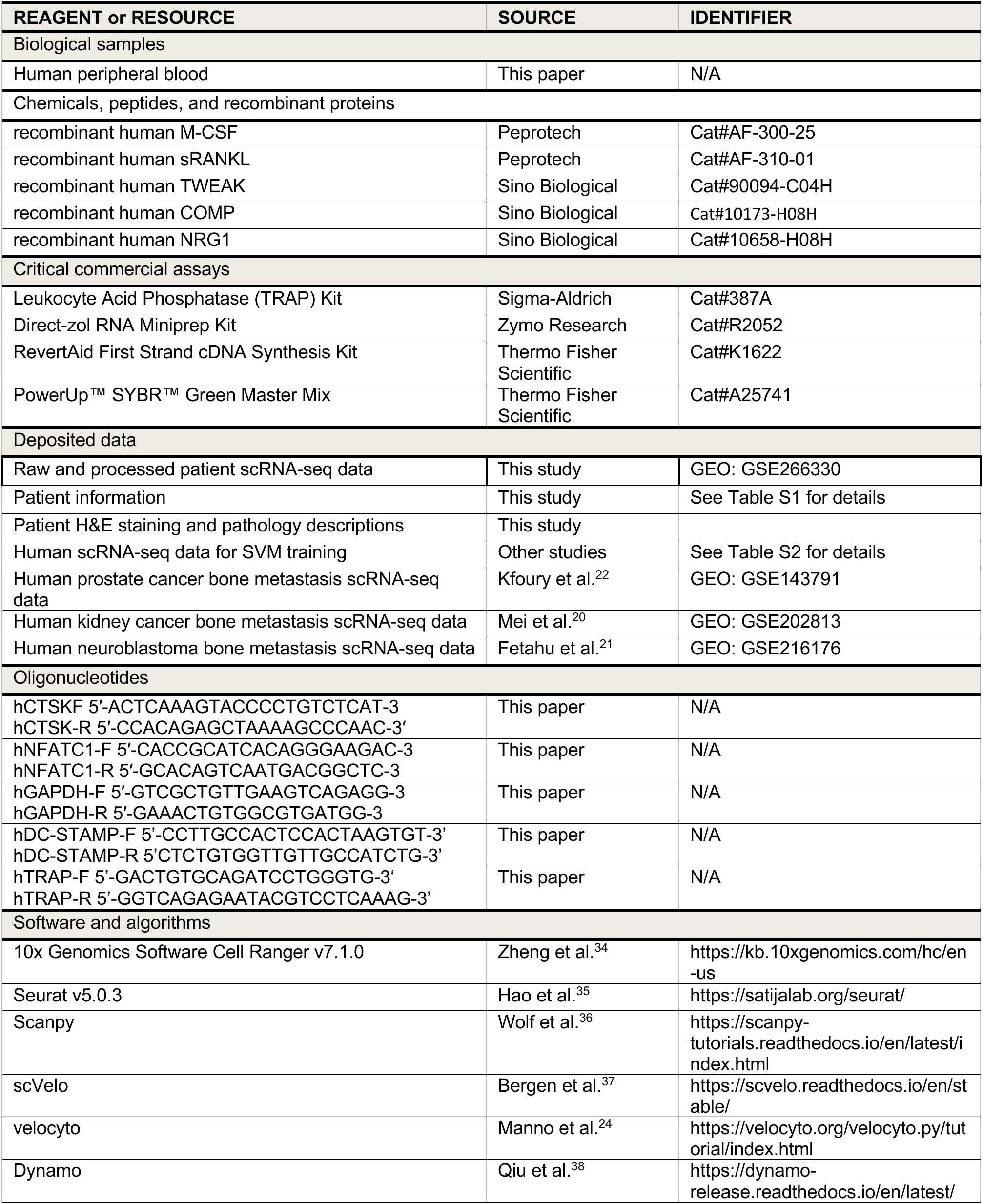
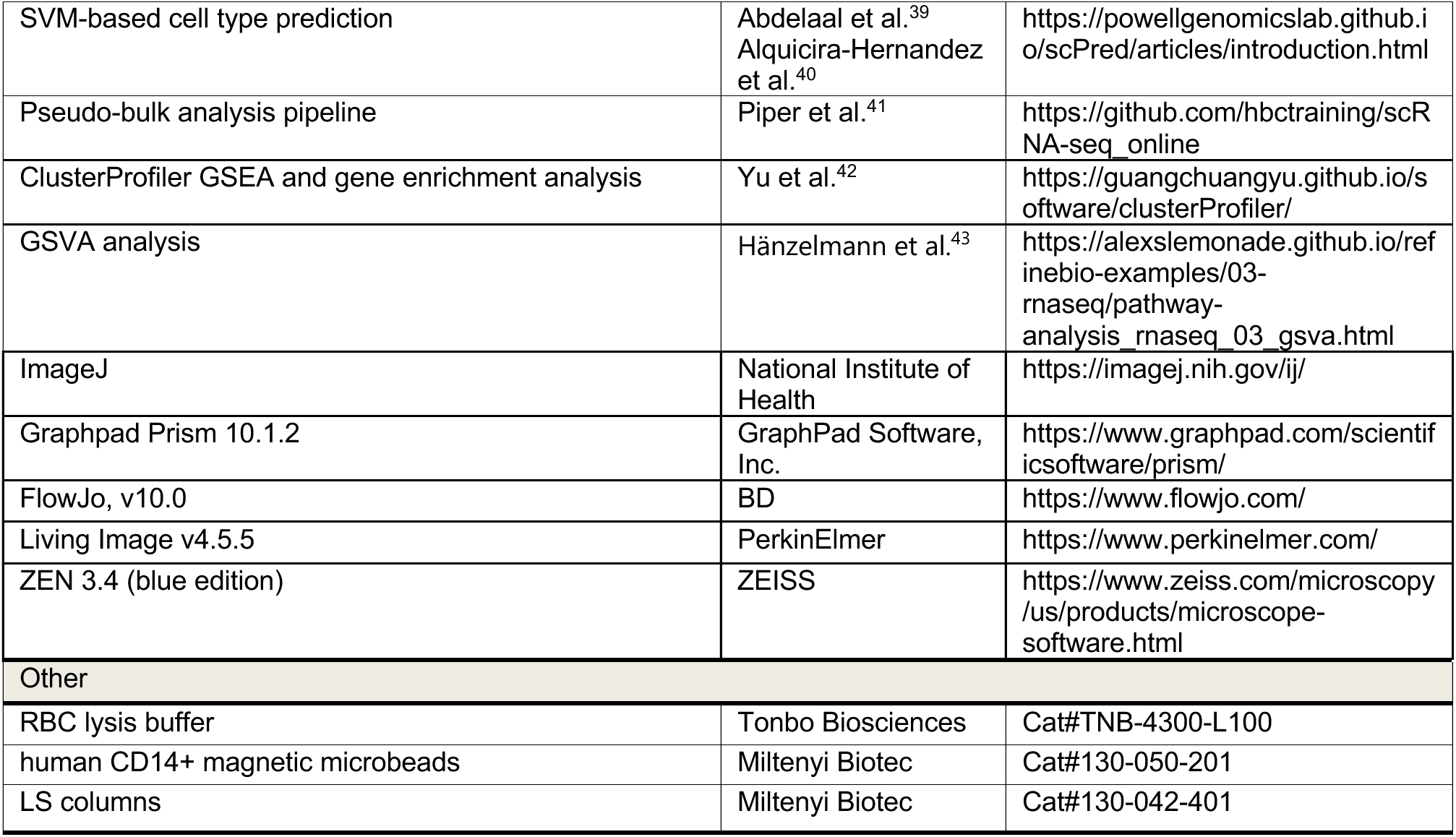

### RESOURCE AVAILABILITY

#### Lead Contact

For additional information or to request resources and reagents, please contact the Lead Contact, Xiang H.-F. Zhang, at.

#### Materials Availability

No new unique reagents were generated in this study.

#### Data and Code Availability

Raw and processed scRNA-seq data has been deposited in the NCBI GEO database (accession number: GSE266330). The code is available upon request.

#### Supplemental information

Document S1 presents high-resolution H&E staining images for a random selection of patients, accompanied by their pathological descriptions.

Table S1 lists patient IDs along with corresponding treatment details, gender, cancer types, and cell counts from our single-cell RNA sequencing dataset.

Table S2 details the datasets utilized to train the SVM for cell type annotations.

Table S3 outlines the selected ligand-receptor pairs identified in the cell-cell communication analysis, specifying the sender and receiver cells involved.

### EXPERIMENTAL MODEL AND SUBJECT DETAILS

#### Patient and healthy samples

This study collected patient samples from MD Anderson Cancer Center in collaboration with Dr. Robert Satcher. The protocols for the collection and use of human bone metastasis samples were performed following the Declaration of Helsinki and approved by the Institutional Review Boards at Baylor College of Medicine (H-49396), The University of Texas MD Anderson Cancer Center (PA15-0225), and the University of Texas Medical Branch (H-46675). All the patients have provided written informed consent on the use of their samples for research purposes when undergoing orthopedic surgery. The selection of patient samples was driven by clinical necessity, focusing on individuals requiring surgical intervention. However, there were no restrictions on patient selection beyond this clinical consideration. Our sample collection was comprehensive, encompassing both single-cell sequencing and the generation of tissue sections for pathology studies as dictated by clinical need. Specifically, we acquired bone tissue regions from 34 patients, encompassing both metastasized tumors and bone tissues. These samples were predominantly sourced from long bones within the human body. Additionally, we obtained 5 healthy individual human bone marrow from Lonza (Catalog#: 1M-105). This approach allowed us to acquire a diverse range of samples tailored to the specific requirements of the patients, ultimately contributing to the comprehensiveness of our study.

### METHOD DETAILS

#### Single-cell RNA library preparation and sequencing of freeze-thawed tissue samples

To optimize cost-effectiveness, we implemented a cell hashing strategy for processing and sequencing the patient samples collected over the years. Fresh tissues, ranging from 0.5 to 2 cm³, were finely minced into 1 mm³ pieces using scalpels and pestles in 24-well plates with 5 ml of DMEM. These tissues were then preserved in liquid nitrogen in a cell-stocking buffer comprising 10% DMSO mixed with 90% FBS in DMEM.

For each batch of samples, typically containing 3-4 specimens, we initiated the process by thawing the samples and washing with PBS. The samples were digested in a dissociation solution containing 1 mg/ml of Collagenase II (Gibco, 17101-015), and 3 mg/ml of Dispase II (Roche, 13437500) in DMEM F12/HEPES medium (Gibco, 113300). In cases where hard bone tissues were present, mechanical grinding of the tissue was conducted before dissociation. The samples were incubated at 37°C in for 30 minutes. Subsequently, the tissue suspension was filtered through a 70-μm strainer to ensure complete tissue grinding, and the strainer was rinsed and ground with DMEM to capture any remaining single cells. The flow-through was then centrifuged at 450g for 5 minutes, and the supernatant was carefully removed. In instances where red blood cells (RBCs) were present in the pellet, 3–4 ml of 1×RBC lysis buffer (1:10 dilution of 10× RBC lysis buffer (MACS, 130-094-183) in Milli-Q water) was applied, with incubation on ice for 10 minutes. To halt and remove RBC lysis, samples were centrifuged at 450g for 5 minutes. The supernatant was discarded, and the cell pellet was washed with 3 ml of 4°C DMEM. After centrifuging at 450g for 5 minutes, the supernatant was discarded, and the cells were resuspended in a cold PBS solution (Sigma, D8537) containing 2% FBS (Gibco, 52567-01). To further eliminate dead cells and cell debris, single cell solution was sorted through FACS sorting machine of DAPI (Invitrogen, R37606) negative, and Ghost dye (Tonbo, 13-0868-T100) negative group. The subsequent steps encompassed single-cell capture, barcoding, and library preparation, following the Chromium Next GEM Single Cell 3ʹ v3.1: Cell Multiplexing protocol (CG000383) and carried out with the assistance of the Single Cell Genomics Core at BCM. The final libraries, containing barcoded single-cell transcriptomes, were sequenced to a depth of 600 million (for transcripts) and 150 million (for cell barcodes) reads on the Novoseq 6000 system (Illumina) at the Genomic and RNA Profiling Core (GARP). Data processing was performed using the CASAVA 1.8.1 pipeline (Illumina), and sequence reads were converted into FASTQ files, cell CMO (Cell Multiplexing Oligo) barcoding matrices, and UMI read counts using the CellRanger software (10X Genomics).

#### Data preprocessing, quality control, and batch correction

We specifically used cellranger multi pipeline to process the multiplexed raw data. For each dataset and patient, we generated multiplexing and count assays based on CellRanger outputs, and all subsequent processing was conducted using Seurat^44^ packages (v.4.0 and v.5.0).

Initially, we performed hashtag demultiplexing within the Seurat framework. To ensure data quality, a two-step doublet filtering process was applied. First, cells were filtered based on the demultiplexing results and hashtag assignments, retaining only the singlets (cells that were identified to have single barcode labeling). Subsequently, cells with fewer than 200 detected genes or exhibiting a deviation of more than two-fold from the median gene count were excluded. Additionally, cells with more than 10% mitochondrial genes were systematically filtered out to minimize mitochondrial gene contamination. We proceeded to normalize each individual dataset using the default parameters of the NormalizeData function. Following normalization, we utilized the FindVariableFeatures function with its default settings to identify variable genes within each normalized dataset. For integration purposes, we employed the SelectIntegrationFeatures function, selecting the most 2,000 variable genes. To identify anchors across all datasets, we compared various methods, including “rpca,” “cca,” and “harmony.” We chose the “rpca” method due to its superior speed and efficiency^44^. These anchors were then used to integrate the matrices using the IntegrateData function, specifying the parameter dims as 1:50. Under the integrated assay, we performed scaling and PCA analysis. Finally, we utilized UMAP for dimension reduction and visualizing the unsupervised clustering results of the data.

#### Machine learning based cell type/state identification

We initially categorized the major cell groups (immune, epithelium, and stromal cells) based on the expression levels of conventional cell markers (immune: PTPRC/CD45 for pan-immune cells; CD3E/G for T cells; CD79A for B cells; CD68 for myeloid cells; EPCAM, KRT8/18/19 for epithelia cells; and PECAM, ACTA2, MYL9 and MYLK for stromal cells). For immune cell annotations, we employed machine learning support vector machines (SVM)^39,40^, we integrated, well-annotated single-cell datasets from various publications: (GSE179346^45^; E-MTAB-8884, E-MTAB-9139 (https://www.ebi.ac.uk/arrayexpress/), GSE175604^46^, GSE120221^47^, GSE193138^48^, GSE159929^49^, GSE181989^50^, GSE159624^51^, GSE130430^52^, GSE139369^53^, GSE165645^54^, GSE128639^55^, GSE185381^56^, GSE166895^57^, GSE133181^58^, GSE135194^59^, resulting in a reference dataset comprising over 670,000 immune cells and encompassing more than 40 cell types and cellular states (**Table S2**). Following cell type predictions in our dataset using the trained references, we conducted manual validation of classical cell marker expression^60^. In the case of stromal cell populations, we primarily utilized a comprehensive single-cell study on bone stromal cells as our reference^61^.

To reinforce our annotations, we employed cell type marker panels sourced from the interactive web portal ToppCell^61^ (http://toppcell.cchmc.org/) for cross-validation purposes. To illustrate the results of this manual validation process, we computed the average expression levels of each reference gene set across all clusters within single-cell datasets, with the same resolution that was utilized for cell type predictions. ^21,22,62^

#### Epithelium copy number variation estimation and malignancy quantification

We utilized the infercnv^25^ R package to investigate subsets of epithelial cells. Notably, since normal bone tissues do not contain epithelial cells, the presence of epithelial cells in our dataset signifies their cancerous nature. To assess copy number changes in the epithelial cells (observation cells), we employed stromal cells and immune cells as reference cells. To quantitively describe the malignancy of epithelial cells, we established a scoring system: assigning 2 marks for each allele with two or more copy number changes (including both addition and deletion), 1 mark for one, and 0 for neutral changes. This allowed us to compute the total scores for each of the infercnv subclusters. High malignant clusters were identified as those subclusters with a median score greater than the 75th percentile of the reference group, while the remaining clusters were classified as low malignant clusters. We then created the epithelial cell malignancy group in our dataset according to such classifications.

#### Determination of patient immunophenotypes

The primary drivers of three distinct immunophenotypes among cancer patients are the varying fractions of different cell types. To uncover these patterns, we used an unbiased hierarchical clustering approach, utilizing the average linkage criterion. This method firstly grouped patients based on cluster fractions (cell number of each cluster divided by the total cell numbers, per patient) within each patient, the clusters are derived from previous unsupervised transcriptomic clustering. Subsequently, we assigned their cellular identities based on pre-determined cell type identities. The cell transcriptomic clusters employed for this analysis were generated at a high resolution (resolution = 5.0) from FindClusters function, and we scaled (z score) the cell fractions within each cluster across rows (patients). To validate our findings, we verified the hierarchical clustering results by selectively plotting the fractions of designated cell types across the patient cohort (supervised clustering).

#### Trajectory inference

We primarily utilized the Dynamo^38^ package and cross validated the results in scVelo^24,37,63^, which leverages RNA velocity to conduct trajectory inference. Our primary focus was on myeloid cells, CD4+ T cells, and CD8+ T cells. We firstly generated loom files from the CellRanger outputs, and then applied these files along with the desired cell type-specific dataset for trajectory analysis. One notable advantage of the Dynamo package is its ability to perform unbiased trajectory analysis without the need for specific “root” assignments. This feature is particularly valuable as it ensures a more objective analysis. Moreover, the trajectory analysis provided compelling evidence for the high accuracy of our cell type annotations. It is important to highlight that cell type annotation and trajectory inference Z-score are distinct processes. However, when we combine the results of trajectory inference with our cell type annotations, they synergistically yield biologically meaningful insights. For instance, the trajectory analysis successfully established connections between precursor cell types and mature cell types, thus elucidating the fulfillment of cellular differentiation paths. Furthermore, we present the output from the state_path function, as it not only reveals estimated developmental routes but also highlights preferred paths in different immunophenotypes, based on the thickness of the arrows. To ensure the robustness of our findings, we further validated our results using alternative packages such as scVelo^24^ and Monocle 3^64^.

#### Cell-cell communication analysis

We conducted cell-cell communication analysis using CellChat^65^ to compare the two major immunophenotypes. Our objective was to identify primary communication senders and receivers, and significant different signaling pathways, focusing on the Mac-OC and Treg-Tex groups compared to healthy donors. To evaluate interaction strength differences between these immunophenotypes for various cell type pairs, we systematically assessed signaling pathways present in at least one group, totaling 190 out of 293 pathways in the CellChat database. We then created an integrated scatter plot to visualize the biased signaling pathways within each immunophenotype group across the entire signaling pathway landscape. The process involved calculating differences in communication probabilities between the two groups for each cell pair within each signaling pathway. Subsequently, we standardized these differences by scaling them as percentages relative to the range for each specific signaling pathway. This analysis allowed us to compile a data matrix for generating scatter plots, distinguishing immunophenotype-specific signaling pathways based on a 75% threshold. Signaling pathways with a positive relative difference above 75% were considered Mac-OC specific, while those with a negative relative difference below −75% were identified as Treg-Tex biased.

#### Transcriptomic differences across cancers and immunophenotypes

We employed PCA to assess transcriptomic variations within our dataset, categorizing cells either by cancer types or immunophenotypes. The variance was computed based on the distribution of each cell type, grouped according to their respective cancer types or immunophenotypes.

#### Pseudo-bulk transcriptomic comparisons and gene enrichment analysis

We opted for pseudo-bulk analysis to compute differentially expressed genes (DEGs) since our aim was to assess cell type-specific expression differences at the population level. We followed these key steps:

- First, we selected cells of interest corresponding to the desired cell type(s) before initiating the differential expression (DE) analysis.
- Next, we obtained raw counts by applying quality control (QC) filtering to the designated cells intended for DE analysis.
- We then integrated the counts and accompanying metadata at the sample level.
- Subsequently, we performed the DE analysis, by using DESeq2^66^ packages in R.

Following the acquisition of DEGs, we conducted Gene Set Enrichment Analysis (GSEA) using the ClusterProfiler^42^ R package. The pathway dataset utilized for this analysis was sourced from MSigDB (https://www.gsea-msigdb.org/gsea/msigdb/). To select top significant differentially regulated pathways, we set up parameters with a stringent BH-adjusted p-value cutoff of 0.01. To visualize the differences in pathway activity across different cell types, cancer types or immunophenotypes, we utilized the AUCell^67^ R package to compute pathway activity scores. Visualization of these scores was achieved through violin plots using the AddModuleScore function. Significance statistics were calculated using the Wilcoxon rank-sum test.

#### Paraffin Embedding, Microtomy and Hematoxylin and Eosin Staining (H&E)

The bone metastasis samples were first fixed in 10% neutrally buffered formalin for up to 24 hours at 4 degrees Celsius. They were then subjected to decalcification in 14% EDTA (pH 7.4) for about one week, with the decalcifying medium changed every 2-3 days. Subsequently, the tissues were delicately transferred into histology cassettes using tweezers and incubated in 70% ethanol before being sent to the pathology core at the breast center of BCM. There, the tissues underwent automated dehydration using ethanol, followed by clearing with xylenes and infiltration with paraffin using the state-of-the-art Sakura Tissue-Tek VIP Processor. Finally, manual embedding of the bone metastasis tissues was performed using the Sakura Tissue-Tek paraffin embedding center. The tissue blocks were cut into 4µm slices using Richard-Allan HM 315 microtomes. Afterward, the sliced slides underwent H&E staining with the Shandon Varistain 24-4 Automatic Slide Stainer.

#### Diagnostic Consultation for H&E-Stained Human Bone Metastasis Samples

Histological features indicative of invasive carcinomas were discerned on H&E-stained slides of human bone metastasis by Dr. George Miles from the pathology core at the breast center of BCM.

#### Isolation of CD14+ Monocytes from Peripheral Blood for Osteoclastogenesis

The peripheral blood (PB) samples were sourced from healthy donors at the Gulf Coast Regional Blood Center in Houston, collected using lithium heparin vacuum tubes. Red blood cell (RBC) lysis was carried out with RBC lysis buffer (Tonbo Biosciences, Cat#TNB-4300-L100). Subsequently, CD14+ monocytes were isolated from the lysed PB cells using human CD14+ magnetic microbeads (Miltenyi Biotec, Cat#130-050-201) with LS columns (Miltenyi Biotec, Cat#130-042-401). These isolated human CD14+ monocytes were cultured in α-MEM supplemented with 10% FBS and antibiotics. Osteoclastogenesis induction was achieved with 25 ng/ml recombinant human M-CSF (Peprotech,Cat#AF-300-25) and 50 ng/ml recombinant human sRANKL (Peprotech,Cat#AF-310-01) following established protocols^68^. Additional treatment with recombinant human TWEAK protein (Sino Biological, Cat#90094-C04H) was added dose-dependently, as illustrated in the figure. TRAP staining was conducted using the Leukocyte Acid Phosphatase (TRAP) Kit (Sigma-Aldrich, Cat#387A) as per the manufacturer’s instructions. The concentration of molecules for *in vitro* osteoclastogenesis and TWEAK, COMP, and NRP1 treatment was set based on published studies^69–71^.

#### mRNA Extraction and qRT-PCR

To extract RNA, the Direct-zol RNA Miniprep Kit (Zymo Research, R2052) was utilized according to the manufacturer’s guidelines. Following RNA extraction, cDNA synthesis was carried out using the RevertAid First Strand cDNA Synthesis Kit (Thermo Fisher Scientific, K1622). For real-time PCR analysis, PowerUp™ SYBR™ Green Master Mix (Thermo Fisher Scientific, A25741) was employed on a CFX96 Real-Time PCR system (Biorad). Quantitative PCR analysis was performed with primer sequences provided in the Key Resources Table.

